# Unraveling Chamber-specific Differences in Intercalated Disc Ultrastructure and Molecular Organization and Their Impact on Cardiac Conduction

**DOI:** 10.1101/2023.02.13.528369

**Authors:** Heather L. Struckman, Nicolae Moise, D. Ryan King, Andrew Soltisz, Andrew Buxton, Izabella Dunlap, Zhenhui Chen, Przemysław B. Radwański, Seth H. Weinberg, Rengasayee Veeraraghavan

## Abstract

During each heartbeat, the propagation of action potentials through the heart coordinates the contraction of billions of individual cardiomyocytes and is thus, a critical life process. Unsurprisingly, intercalated discs, which are cell-cell contact sites specialized to provide electrical and mechanical coupling between adjacent cardiomyocytes, have been the focus of much investigation. Slowed or disrupted propagation leads to potentially life-threatening arrhythmias in a wide range of pathologies, where intercalated disc remodeling is a common finding. Hence, the importance and urgency of understanding intercalated disc structure and its influence on action potential propagation. Surprisingly, however, conventional modeling approaches cannot predict changes in propagation elicited by perturbations that alter intercalated disc ultrastructure or molecular organization, owing to lack of quantitative structural data at subcellular through nano scales. In order to address this critical gap in knowledge, we sought to quantify intercalated disc structure at these finer spatial scales in the healthy adult mouse heart and relate them to function in a chamber-specific manner as a precursor to understanding the impacts of pathological intercalated disc remodeling. Using super-resolution light microscopy, electron microscopy, and computational image analysis, we provide here the first ever systematic, multiscale quantification of intercalated disc ultrastructure and molecular organization. By incorporating these data into a rule-based model of cardiac tissue with realistic intercalated disc structure, and comparing model predictions of electrical propagation with experimental measures of conduction velocity, we reveal that atrial intercalated discs can support faster conduction than their ventricular counterparts, which is normally masked by inter-chamber differences in myocyte geometry. Further, we identify key ultrastructural and molecular organization features underpinning the ability of atrial intercalated discs to support faster conduction. These data provide the first stepping stone to elucidating chamber-specific impacts of pathological intercalated disc remodeling, as occurs in many arrhythmic diseases.

## INTRODUCTION

Fast and reliable propagation of action potentials through the myocardium is a critical life process, the disruption of which can lead to life-threatening arrhythmias. Structural differences between atrial and ventricular myocardium at the organ, tissue, and cellular levels are well established determinants of differences in action potential propagation under normal physiological conditions^1, 2^, as well as determining chamber-specific arrhythmia susceptibility under pathological conditions. Intercalated discs (IDs), which are cell-cell contact sites specialized to provide electrical and mechanical coupling between adjacent cardiomyocytes, are thus a key focus of research into cardiac action potential propagation with a growing body of evidence highlighting the strong influence of ion channel localization to specialized ID nanodomains^3–7^. However, there is limited data on ID ultrastructural and molecular organization at subcellular through nano scales, while mounting evidence highlights their importance in determining function^3–8^. Additionally, conventional modeling approaches cannot predict conduction changes in response to perturbations that alter ID ultrastructure or molecular organization. Quantifying structure-function relationships at these finer spatial scales and in a chamber-specific manner is crucial to understanding cardiac action potential propagation during normal physiological conditions and its modes of failure under pathological perturbation, and thereby, developing effective therapies for the prevention of arrhythmias. Thus, we tested the hypothesis that chamber-specific heterogeneities in conduction and its stability under perturbation are influenced by underlying heterogeneities in ID ultrastructure and molecular organization.

To this end, we applied a combined experimental and modeling approach to compare the structural basis of conduction between healthy adult murine atria and ventricles. First, we systematically quantified myocyte geometry, ID ultrastructure, and molecular organization using light and electron microscopy. Next, we incorporated these measurements into a rule-based model of cardiac tissue with realistic ID structure, and compared model predictions of electrical propagation with experimental measures of conduction velocity from optical mapping of *ex vivo* hearts. Consistent with previous findings, action potential propagation was faster in ventricular myocardium than in atrial myocardium. However, model predictions revealed that the conduction differences under normal physiological conditions were largely a reflection of differences in cell geometry, and not ID characteristics. Counterintuitively, after accounting for cell size differences between chambers, the ultrastructure and molecular organization of atrial IDs were found to support faster conduction, compared to ventricular IDs. In other words, differences in cell size mask underlying heterogeneities in ID structure, which may be unmasked by pathological remodeling with important implications for the risk of proarrhythmic conduction defects.

## MATERIALS AND METHODS

All animal procedures were approved by The Ohio State University Institutional Animal Care and Use Committee and performed in accordance with the Guide for the Care and Use of Laboratory Animals published by the U.S. National Institutes of Health (NIH Publication No. 85-23, revised 2011).

### Tissue Collection and Myocyte Isolation

Adult, male (aged 12-34 weeks) wild-type (WT) mice (C57BL6 background), purchased from Jackson Laboratories (Cat. #000664), were anesthetized with 5% isoflurane + 100% oxygen (1L/min). Once the mice were unconscious, anesthesia was maintained within 3-5% isoflurane + 100% oxygen (1L/min). Once the animal was in a surgical plane of anesthesia, the heart was excised and submerged in cold electrolyte solution (containing in mM: NaCl 140, KCl 5.4, MgCl_2_ 0.5, dextrose 5.6, HEPES 10; pH adjusted to 7.4) to remove any remaining blood prior to myocyte isolation, cryopreservation (for immunofluorescence), fixation (for transmission electron microscopy [TEM]), or Langendorff perfusion (for optical mapping) as previously described^8–10^.

For light microscopy studies in tissue, hearts were cryopreserved in OCT compound (Fisher Health Care, Houston, TX, USA) using vapor phase nitrogen to ensure uniform freezing. For TEM, ∼0.5mm cubes of tissue were fixed overnight in 2.5% glutaraldehyde followed by a 5 min wash in PBS and storage in PBS for further processing. For isolated myocyte studies, Langendorff-perfused hearts were enzymatically digested using Liberase™ TH (Roche, Basel, Switzerland) enzyme in Ca^2+^-free electrolyte solution. After digestion, the atria and ventricles were dissected and separately minced in electrolyte solution containing 2% Bovine Serum Albumin (BSA; Sigma-Aldrich, Bornem, Belgium), dispersed by gentle agitation, and filtered through a nylon mesh. Cardiomyocytes are resuspended in low Ca^2+^ electrolyte solution (containing 0.1 mM CaCl_2_). Then, cells were plated on laminin-coated glass coverslips, fixed for 5 min with 2% paraformaldehyde (PFA), washed in phosphate-buffered saline (PBS) (3 × 10 min) and stored in PBS at 4°C for immunolabeling.

All structural imaging (light and electron) was performed on the left atrium and left ventricular free wall to focus on working myocardium.

### Fluorescent Immunolabeling

Tissue was cryosectioned (5 μm thickness) and fixed with 2% PFA (5 min at room temperature [RT]) followed by a PBS wash (3 x 10 min at RT). Samples (either fixed tissues or cells) were permeabilized with 0.2% Triton X-100 in PBS (15 min at RT) and incubated with blocking agent (1% bovine serum albumin, 0.1% triton in PBS for 2 hrs at RT). For sub-diffraction confocal imaging (sDCI), samples were labeled with primary antibodies (overnight at 4°C) and washed in PBS (3 x 5 min at RT) prior to labeling with secondary antibodies (2 hrs at RT). After secondaries, samples were washed in PBS (3 x 5 min at RT), mounted in Prolong Gold (Invitrogen by Thermo Fisher Scientific, Grand Island, NY, USA) and allowed to cure for 48 hours at room temperature before imaging. A similar protocol was used to prepare samples for STochastic Optical Reconstruction Microscopy (STORM) with the following minor changes. A 10x diluted solution of the blocking agent (.1% bovine serum albumin, 0.05% triton in PBS) was used for washes before and after secondary antibodies in place of PBS. Instead of being mounted on glass slides, labeled samples on 20 mm coverslips thickness #1.5 (Electron Microscopy Sciences, Hatfield, PA, USA) were post-fixed in 2% PFA (5 minutes at RT) followed by PBS washes (3 x 5 min at RT), then optically cleared in Scale U2 buffer (4M urea + 30% glycerol + 0.1% triton in water at 4°C) for 48 hours before imaging.

Proteins of interest were labeled with well-validated custom and commercial antibodies: Connexin43 (Cx43; mouse monoclonal antibody, Cat. #: MAB3067, EMD Millipore Corp., Darmstadt, Germany; rabbit polyclonal antibody, Cat. # C6219, Millipore Sigma, St. Louis, MO), N-cadherin (N-cad; mouse monoclonal antibody, Cat. # 610921, BD Biosciences, San Jose, CA), desmoplakin (Dsp; rabbit polyclonal antibody, Cat. # ab71690, Abcam, Waltham, MA), inward-rectifier potassium channel 2.1 (K_ir_2.1; rabbit polyclonal antibody, Cat. # APC-159, Alomone labs, Jerusalem, Israel), cardiac voltage-gated sodium channel 1.5 (Na_V_1.5; a validated custom rabbit polyclonal antibody^4^, and sodium/potassium ATPase (NKA; a previously validated custom monoclonal antibody^11^. Samples labeled for confocal microscopy were labeled with goat anti-mouse and goat anti-rabbit secondary antibodies conjugated to Alexa 488 and Alexa 568 were used (1:4000; ThermoFisher Scientific, Grand Island, NY). For STORM, samples were labeled with goat anti-mouse Alexa 647 (1:1000) and goat anti-rabbit Biotium CF 568 (1:2000) secondary antibodies (ThermoFisher Scientific, Grand Island, NY, USA). Simultaneous labeling with two primary antibodies from the same animal background was accomplished by completing the staining protocol with one set of primary and secondary antibodies followed by blocking with a donor secondary antibody (conjugated to fluorophore to assess degree of blocking) to block any remaining unbound 1^st^ antibodies. A second round of labeling was performed with a second combination of primary and secondary antibodies.

### Confocal Imaging

Confocal imaging was performed using an A1R-HD laser-scanning confocal microscope equipped with four solid-state lasers (405, 488, 560, and 640 nm, 30 mW each), a 63×/1.4 numerical aperture oil immersion objective, two GaAsP detectors, and two high sensitivity photomultiplier tube detectors (Nikon, Melville, NY, USA). Images were collected as single plane or z-stacks with sequential spectral imaging (line-wise) in order to avoid spectral mixing. In addition, images were collected with Nyquist sampling (or greater) and post-processed with 3D deconvolution, as previously described^12^. Spatial analysis of images was performed to quantify signal abundance and localization (distance from fluorescent signals to another signal or a cellular landmark structure) using morphological object localization (MOL), a custom algorithm implemented in Matlab (Mathworks Inc, Natick, MA)^13^.

### Transmission Electron Microscopy (TEM)

After 2.5% glutaraldehyde fixation (SIGMA-ALDRICH, Saint Louis, MO, USA) as previously described in the tissue collection section, samples were postfixed with 1% osmium tetroxide (Ted Pella Inc., Redding, CA, USA) and then *en bloc* stained with 1% aqueous uranyl acetate (Ted Pella Inc., Redding, CA, USA), dehydrated in a graded series of ethanol, and embedded in Eponate 12 epoxy resin (Ted Pella Inc., Redding, CA, USA). Ultrathin sections (70 nm) were cut with a Leica EM UC7 ultramicrotome (Leica microsystems Inc., Deerfield, IL), collected on copper grids, and then stained with Reynold’s lead citrate and 2% uranyl acetate. Images were collected using a FEI Tecnai G2 Spirit transmission electron microscope (Thermo Fisher Scientific, Waltham, MA) operating at 80kV, and a Macrofire (Optronics, Inc., Chelmsford, MA) digital camera and AMT image capture software.

In order to assess multiscale structural properties of IDs, paired images of each ID were collected at 6,000x, 10,000x, and 20,000x magnifications. Morphometric quantification was performed using ImageJ (National Institutes of Health, http://rsbwe b.nih.gov/ij/) by manual identification and quantification of the following 21 specific measurements. Color coded annotations of each ID measurement are overlaid on TEM images at each magnification in figures 3E-F, 4H-I, 5F, 6C-E. Images at 6,000x magnification were used to quantify total ID length (cross-sectional length) (Figure 3E), and length of the plicate and interplicate subdomain regions (Figure 3F). Images at 10,000x magnification were used to quantify the frequency and amplitude of plicate membrane folds (Figure 4H), and the lengths of MJs and GJs in both plicate and interplicate regions (Figure 4H, 5F). Images at 20,000x magnification were used to quantify the intermembrane distance at multiple sites within/near MJs and GJs (Figure 4I, 5F). Ratios of these measurements across provided further insight into ID structural organization (Supplementary Table 1).

### Super-Resolution Microscopy

STochastic Optical Reconstruction Microscopy (STORM) was performed using a Vutara 352 microscope (Bruker Inc, Middleton, WI, USA) equipped with biplane three-dimensional detection, and fast scientific complementary metal–oxide–semiconductor (sCMOS) camera achieving 20 nm lateral and <50 nm axial resolution. Individual fluorophore molecules were localized with a precision of 10 nm. Registration of the two color channels was accomplished using localized positions of several TetraSpeck Fluorescent Microspheres (ThermoFisher Scientific, Carlsbad, CA, USA) scattered throughout the field of view. 3-dimensional images of *en face* IDs were collected at 150 frames/z-step. Machine learning-based cluster analysis of single-molecule localization data was performed using STORM-based relative localization analysis (STORM-RLA), as previously described^14^.

### Optical Mapping

Optical mapping of Langendorff-perfused hearts was performed as in previous studies^4, 15^. Briefly, murine hearts were perfused as Langendorff preparations with an electrolyte solution containing (in mM): 1.8 CaCl_2_·2H_2_O, 143 NaCl, 4.2 KCl, 5.5 Dextrose, 1 MgCl_2_·6H_2_O, 1.2 NaH_2_PO_4_, 5.5 NaOH (pH 7.42). Temperature was maintained at 36°C and hearts were electrically stimulated with an AgCl_2_ pacing electrode placed septally on the anterior epicardium.

Hearts were perfused with the voltage-sensitive fluorophore, di-4-ANEPPS (7.5 µM, Biotium, CA) and positioned to center the anterior descending artery within the field of view. The anterior epicardium was mapped with a MiCAM03-N256 CMOS camera (SciMedia: 256×256 pixels, field of view 11.0 x 11.0 mm, 1.8kHz frame rate). Motion was reduced with the electromechanical uncoupler blebbistatin (10 µM). For atrial maps, the atria were cut along the septum on the posterior wall and pinned to a sylgard plate for imaging.

Optical signals were recorded at a cycle length of 100ms and conduction velocity (CV) calculated using BV Workbench software (Version 2.7.2). Transverse conduction velocity (CV_T_) is reported as an average of local CV vectors along the short-axis of cardiac tissue. Longitudinal CV (CV_L_) is reported as an average of local CV vectors along the long-axis of cardiac tissue. Conduction velocities were measured from the left atrium and left ventricular free wall.

### Statistical Analysis

The two-sample Kolmogorov-Smirnov test was used to compare whole distributions of measurements between two samples. The Wilcoxon rank sum test was used to compare median values between two populations for non-normal data. A paired two sample Student’s T-test was used to compare means between normally distributed data. A p value ≤ 0.05 was considered statistically significant. Varying degrees of significance are denoted in figures: * for ≤0.05, ** for ≤0.01, *** for ≤0.001, **** for ≤0.0001. All measurements are reported as median, mean ± SEM, quantity of measurements (N), and the first and third quartile range.

### Finite Element Modeling (Ultrastructure integration & Molecular integration)

Cardiac tissue simulations incorporating the effects of atrial- or ventricular-specific ID nanoscale structure and ion channel organization were based on our recently developed computational model formulation^9^, with two key additional features (described below). In brief, we used a ‘rule-based’ approach to construct a 3-D finite element mesh of the ID and the intercellular cleft space (i.e., space between adjacent myocytes) that reproduce key ID measurements, specifically: ID diameter, plicate and interplicate region length, plicate and interplicate region intermembrane distance, GJ size and distribution in both regions, and plicate fold amplitude and frequency. Structural features (e.g., GJ clusters) were generated using a ‘map’ generation algorithm that reproduces the position of individual clusters within the finite element mesh, with the parameters of the algorithm fitted to match the corresponding TEM measurements (e.g., GJ size and density, for each ID region). Following this rule-based approach, individual ID components were constructed into a complete mesh of the cleft space, incorporating several orders of magnitude in detail, from intermembrane separation (nanoscale), up to interplicate and plicate regions (microscale).

While this finite element mesh model of the ID and cleft space captures high-resolution structural detail, incorporation of this mesh into a tissue scale model is not computationally feasible at present. Therefore, we calculated an equivalent electrical network for the resistance within and out of the intercellular cleft space based on the finite element mesh. We calculated this reduced network by partitioning the full mesh into a tractable number (100) of compartments and solving the full electrostatics problem (i.e., Laplace’s equation) on each pair of adjacent compartments, for which the solution enables calculation of the equivalent electrical resistance that in turn define the reduced electrical network. Finally, we incorporated this cleft network into a cardiac tissue model, in which neighboring myocytes are coupled via both GJs and their shared intercellular cleft space. The full model description, equations, parameters, and numerical methods are provided in our recent work.^9^

In our previous simulations, we assumed that all ion channels are uniformly distributed within the ID, such that the conductance of the corresponding ionic currents in each mesh partition is defined as proportional to the partition membrane surface area. Here, we modeled the non-uniform distribution at the ID membranes for Na_V_1.5, K_ir_2.1, and NKA. The position of ion channel clusters within the mesh were generated with a similar map algorithm used to generate the mesh GJs^9^. In addition to cluster size and density, an additional map parameter defines the likelihood of a cluster based on its distance relative to the nearest GJ, with all parameters defined independently for the plicate and interplicate regions. The nearest-neighbor distribution for the channel-GJ from the mesh generation was directly compared with the corresponding STORM-based distribution, and the MATLAB genetic algorithm optimization (Mathworks) was used to fit map parameters. After thus fitting each electrogenic protein’s spatial distribution separately, we defined the conductance of each ionic current in each mesh partition as proportional to the channel density in each partition, which directly represents the electrogenic protein distribution in the tissue model. Thus, each mesh partition had a fraction of the overall ID conductance for each channel modeled. The ID localization values were (as a fraction of total cell conductance) 0.7 for Na_V_1.5 and 0.2 for K_ir_2.1 and NKA. Further, the tissue gap junction conductance was scaled by the total GJ area of the generated mesh, with the ventricular GJ area and conductance used as references and atrial GJ conductance was therefore proportional to the fraction between atrial and ventricular total GJ area. Additionally, in our previous simulations, we assumed that only Na^+^ ions varied in the extracellular cleft space. Here, all ions varied depending on local electro-chemical gradients. Specifically, the three main cations (Na^+^, K^+^ and Ca^2+^) and a generic anion (A^-^), included to maintain electroneutrality in the bulk extracellular space, were dynamic and governed by both ID-localized ionic currents and electrodiffusion.

Chamber-specific ionic currents on both the lateral and ID membranes were governed by the ionic models for atrial^16^ and ventricular^17^ tissues, respectively. Tissue models, representing a one-dimensional “chain” of coupled myocytes, were comprised of 50 cells, with either a chamber-specific or a constant cell length. A propagating electrical wave was initiated by stimulating the first cell at a periodic interval of 500 ms. The resulting system of equations was solved by an operator splitting method, in which voltages were solved by an implicit backward Euler method and gating variables and ionic concentrations were solved with the forward Euler method, with a variable time step. Conduction velocity was measured by calculating the propagation time across the middle 50% of the tissue.

## RESULTS

### Cardiomyocyte and Intercalated Disc Dimensions

Macroscopic differences in atrial and ventricular tissue structure, such as pectinate vs. trabecular muscle architecture or differences in wall thickness, are readily apparent (Figure 1A) and their effects on electrophysiology are well established^1, 2^. However, structural differences between atrial and ventricular IDs, which may underlie conduction differences, have yet to be systematically assessed. Therefore, we set out to quantify structural heterogeneities between the atrial and ventricular IDs on cellular, ultrastructural, and molecular scales.

**Figure 1.**
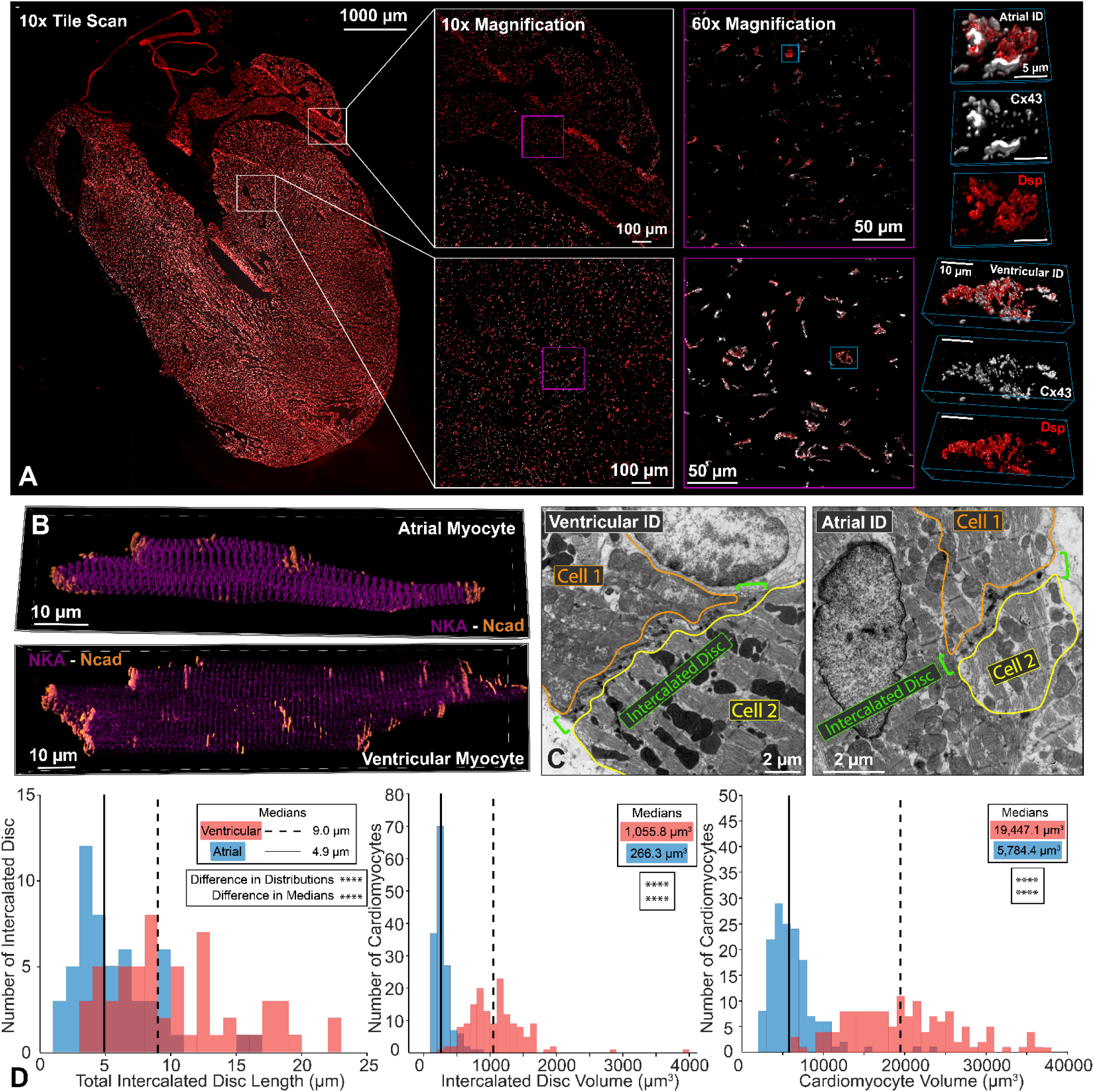
Cardiomyocyte and Intercalated Disc Dimensions. Representative confocal images of atrial and ventricular **A)** intercalated discs from a whole-heart frontal section (gap junctions [Cx43; white] and desmosomes [Dsp; red]) and **B)** isolated cardiomyocytes (adherens junctions [Ncad; orange] and sodium/potassium ATPase [NKA; purple]). Representative thin section **C)** transmission electron microscopy (TEM) images of atrial and ventricular intercalated discs. Intercalated disc size was quantified by **D)** length (left; n= 54[A], 58[V] measurements from 3 hearts) and volume (middle; n= 151[A], 150[V] measurements from 4 hearts). Cardiomyocyte size was quantified by **D)** volume (right; n= 151[A], 150[V] measurements from 4 hearts). Distributions are represented in red for ventricular and blue for atrial intercalated discs. Medians are represented as dashed lines for ventricular and solid lines for atrial results. Differences in distributions and medians were statistically tested by two-sample Kolmogorov-Smirnov test and Wilcoxon signed-rank test, respectively (p > 0.05: ns, p ≤ 0.05: *, p ≤ 0.01: **, p ≤ 0.001: ***, p ≤ 0.0001: ****).

At the cellular scale, we measured differences in cell size and ID dimensions by comparing 3D confocal images of atrial and ventricular cardiomyocytes and tissue. Isolated ventricular cardiomyocytes had larger length, width, depth and volume, compared to atrial myocytes (Figure 1B, D; Supplementary Figure 1A-C; n= 151[A], 150[V] measurements from 4 hearts)^18–22^. Visual inspection of confocal images of IDs from murine cardiac tissue sections suggested that ventricular IDs are substantially larger relative to atrial IDs. (Figure 1A). Therefore, we quantified ID volume from confocal images of isolated cardiomyocytes and cross-sectional length using thin section transmission electron microscopy (Figure 1B-C; volume n= 151[A], 150[V] measurements from 4 hearts; length n= 54[A], 58[V] measurements from 3 hearts). Ventricular IDs had larger volume and cross-sectional length when compared to atrial IDs (Figure 1D). These results demonstrate that atrial and ventricular IDs are structurally disparate and prompt a more thorough investigation of differences across spatial scales.

### Morphological Properties of Intercalated Disc Subdomains

#### ID Structural Landmarks

Next, we characterized ID ultrastructure from micro-through nano-scales via TEM. Within the ID (green brackets; Supplementary Figure 2: left), mechanical coupling is predominantly maintained in the plicate subdomain (magenta box) and electrical coupling in the interplicate subdomain (light blue box; Supplementary Figure 2: middle)^9, 23–30^. It should be noted that small subsets of electrical and mechanical junctions are respectively located within plicate and interplicate regions (Supplementary Figure 3). Mechanical coupling is maintained via adherens junctions (orange box; Supplementary Figure 2: right) and desmosomes (yellow box; Supplementary Figure 2: right), while electrical coupling is provided by gap junctions (red box, Supplementary Figure 2: right). Also localized to juxta-junctional ID nanodomains are electrogenic proteins (black boxes, Supplementary Figure 2: right) crucial for cardiac conduction. Thus, we systematically assessed ID structure at different spatial scales through electron microscopy and used confocal and super-resolution light microscopy to quantify the spatial organization throughout the myocyte and within the ID of ion channels and pumps responsible for the action potential upstroke: the cardiac isoform of the voltage-gated sodium channel (Na_V_1.5; green), inward-rectifier potassium channel (Kir2.1; white), and sodium potassium ATPase (NKA; purple) (Supplementary Figure 2: right). In all these studies, we utilized connexin 43 (Cx43) as a marker for gap junctions at smaller spatial scales and interplicate ID regions at larger scales. Likewise, we utilized N-cadherin (N-cad) and desmoplakin (Dsp) as markers of adherens junctions and desmosomes respectively at smaller scales and of plicate ID regions at larger scales.

#### Plicate and Interplicate Subdomains

At the subdomain level, the ultrastructure of interplicate and plicate regions were characterized in terms of regional length and percent contribution to total ID length (Supplementary Figure 4E-F; length: plicate n= 174[A], 247[V], interplicate n= 97[A], 160[V] measurements from 3 hearts), and (percent: plicate n= 53[A], 58[V], and interplicate n= 53[A], 55[V] measurements from 3 hearts). These ultrastructural features are labeled in colors matched with corresponding labels on the TEM figures. There was no substantial difference in plicate and interplicate length or percent contribution to ID length (Supplementary Figure 4A-D). Plicate and interplicate subdomains were similar in length between chambers. Median plicate length was measured at 1.35 ± 0.78 μm and 1.22 ± 0.47 μm and interplicate median lengths at 1.34 ± 0.51 μm and 1.23 ± 0.74 μm, respectively, for ventricles and atria (Supplementary Figure 4B, D). The percent of the ID comprised of plicate and interplicate subdomains were also similar between chambers (Supplementary Figure 4A, C).

#### Junctional Structures within Plicate Subdomains

Next, we characterized nanodomains located within the plicate and interplicate ID regions. The undulation of the plicate waves was characterized by frequency and amplitude. Ventricular plicate regions had higher amplitude (teal trace) than atrial plicate waves, while there was no difference in frequency (yellow trace) (Figure 2A-B, 2H; amplitude n= 79[A], 128[V], and frequency n= 72[A], 110[V] measurements from 3 hearts)^23, 25, 31^. Although, we observed some distinct desmosomes and adherens junctions, a large proportion of area composita were also observed. Therefore, all mechanical junctions were pooled into a single category for ultrastructural measurements. Plicate mechanical junction length (orange trace) was smaller in atria than ventricular; however, mechanical junctions accounted for the same percent of ID length in both chambers (Figure 2C-D, 2H; length n= 282[A], 329[V], and percent n= 57[A], 52[V] measurements from 3 hearts). These data suggest that atrial IDs contain more numerous, smaller mechanical junctions. Although far less common than mechanical junctions, some gap junctions were observed in the plicate regions. As with mechanical junctions, these were smaller in the atria than in the ventricles but accounted for similar fractions of the plicate membranes in both chambers (Supplementary Figure 3A-B; length n= 29[A], 56[V], and percent n= 19[A], 17[V] measurements from 3 hearts). Thus, these data suggest that plicate gap junctions are smaller yet more numerous in the atria than in the ventricles. The intermembrane distance between the apposed cell membranes was assessed inside mechanical junctions as well as 10 and 30 nm outside (Figure 2I). Intermembrane spacing was similar at all positions relative to the mechanical junctions with atria showing a shift toward smaller values when compared with ventricles (Figure 2E-G; inside n= 178[A], 323[V], 10 nm outside n= 178[A], 241[V], and 30 nm outside n= 178[A], 240[V] measurements from 3 hearts). These data indicate tighter intermembrane spacing within the plicate ID in the atria relative to the ventricles.

**Figure 2.**
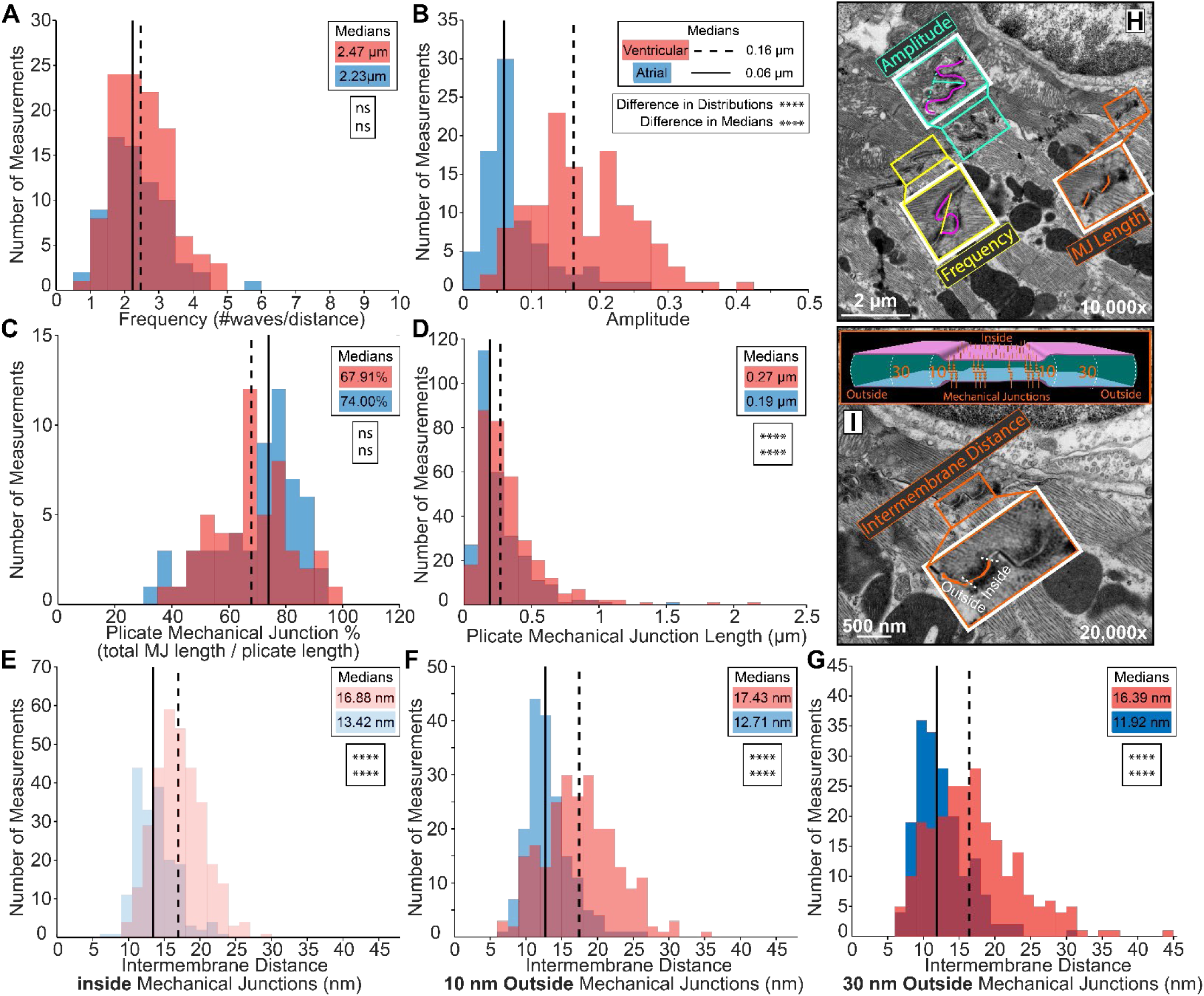
Intercalated Disc Nanodomain Structures in the Plicate Subdomain. Representative TEM images annotated with intercalated disc nanodomain structural measurements within plicate regions: **H)** membrane fold frequency (yellow trace), amplitude (teal trace), mechanical junction length (orange trace), **I)** intermembrane distance inside and outside (10, 30 nm) mechanical junctions (orange fill, cartoon). Plicate fold periodicity was characterized by **A)** frequency (n= 72[A], 110[V] measurements), and **B)** amplitude (n= 79[A], 128[V] measurements). Plicate mechanical junctions were characterized by **C)** percentage relative to the plicate region (n= 57[A], 52[V] measurements), **D)** length(n= 282[A], 329[V] measurements), and **E-G)** intermembrane distance inside and outside (10, 30 nm) mechanical junctions (inside n= 178[A], 323[V], 10 nm outside n= 178[A], 241[V], and 30 nm outside n= 178[A], 240[V] measurements). Performed on 3 hearts. Distributions are represented in red for ventricular and blue for atrial measurements. Medians are represented as dashed lines for ventricular and solid lines for atrial measurements. Differences in distributions and medians were statistically proven with two-sample Kolmogorov-Smirnov test and Wilcoxon signed-rank test, respectively (p > 0.05: ns, p ≤ 0.05: *, p ≤ 0.01: **, p ≤ 0.001: ***, p ≤ 0.0001: ****).

#### Junctional Structure within Interplicate Subdomains

Next, we turned our attention to interplicate regions, where gap junctions were larger and more frequent than in plicate regions (Figure 3). Gap junctions comprised a smaller fraction of interplicate regions (Figure 3A) and were smaller in size in atria than in ventricle (Figure 3B; Figure 3F, red line trace, bottom box; percent n= 40[A], 72[V], and length n= 162[A], 186[V] measurements from 3 hearts). Together, these data suggest that atrial IDs contain fewer, smaller gap junctions resulting in a lower perinexus availability than ventricular IDs. As with gap junctions in the plicate regions, mechanical junctions were rarely found in the interplicate regions. Intriguingly, interplicate mechanical junctions were smaller in the ventricles than in atria but these structures accounted for similar fractions of interplicate regions in both chambers (Supplementary Figure 3C-D; length n= 57[A], 24[V], and percent n= 11[A], 15[V] measurements from 3 hearts). Intermembrane distance was assessed at sites 10, 30, and 50 nm outside gap junctions (Figure 3F). Intermembrane distances increased with increasing distance from gap junctions in both chambers (Figure 3C-E, (10 nm outside n= 58[A], 142[V], 30 nm outside n= 58[A], 143[V], and 50 nm outside n= 58[A], 140[V] measurements from 3 hearts). At 10 nm, intermembrane distances were similar between the chambers (Figure 3C), whereas, at 30 and 50 nm, the ventricles evidenced a larger subpopulation of sites with wider spacing compared to the atria (Figure 3D-E). These data suggest tighter intermembrane spacing within the interplicate ID in the atria relative to the ventricles.

**Figure 3.**
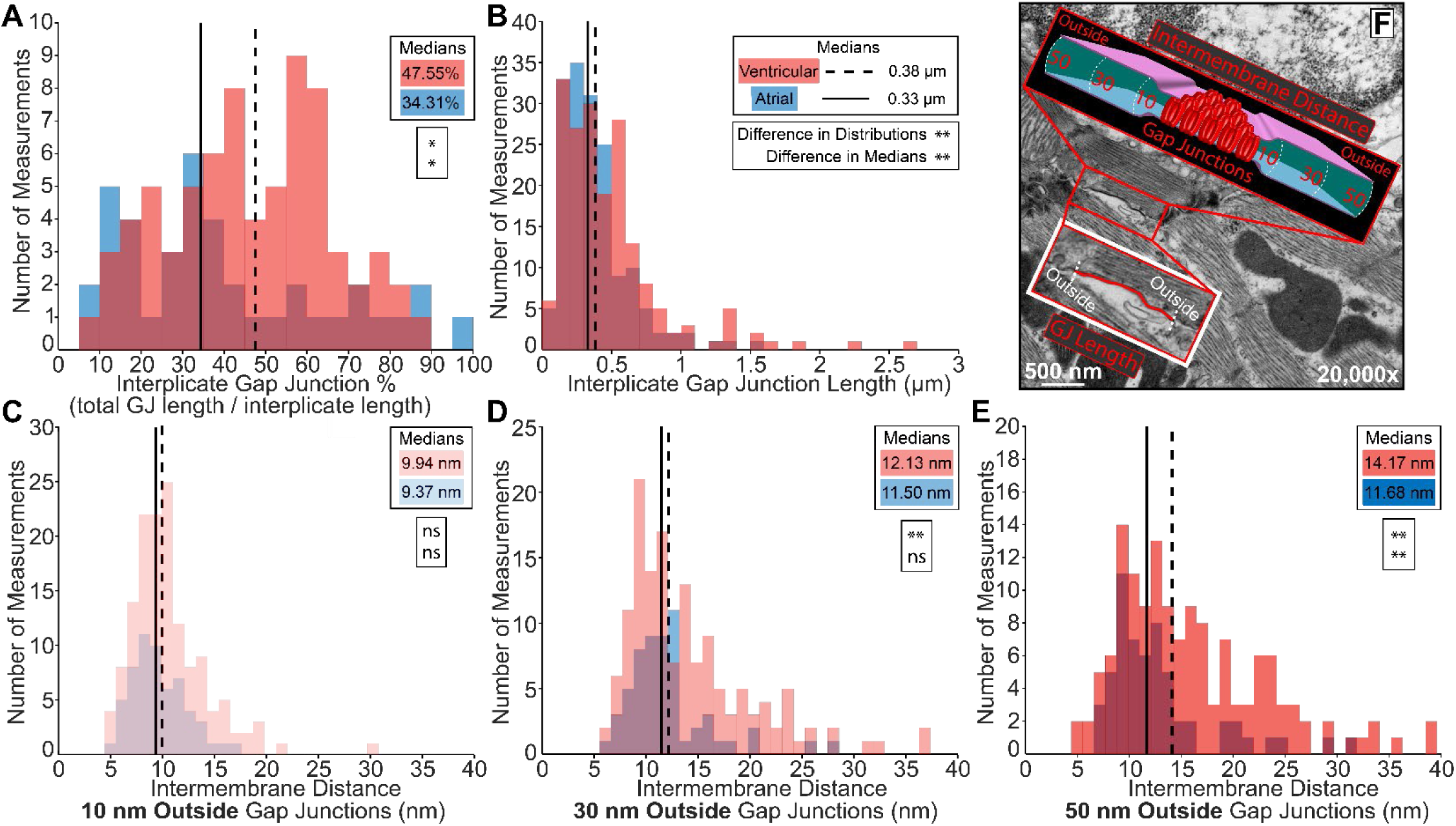
Intercalated Disc Nanodomain Structures in the Interplicate Subdomain. Representative TEM images annotated to show intercalated disc nanodomain structural measurements in the interplicate regions: **F)** gap junction length (red trace), and intermembrane distance outside (10, 30, 50 nm) gap junctions (cartoon). Interplicate gap junctions were characterized by **A)** percentage relative to the interplicate region (n= 40[A], 72[V] measurements), **B)** length (n= 162[A], 186[V] measurements), and **C-E)** intermembrane distance outside (10, 30, 50 nm) gap junctions (10 nm outside n= 58[A], 142[V], 30 nm outside n= 58[A], 143[V], and 50 nm outside n= 58[A], 140[V] measurements). Performed on 3 hearts. Distributions were represented in red for ventricular and blue for atrial measurements. Medians were represented as dashed line for ventricular and solid line for atrial. Differences in distributions and medians were statistically proven with two-sample Kolmogorov-Smirnov test and Wilcoxon signed-rank test, respectively (p > 0.05: ns, p ≤ 0.05: *, p ≤ 0.01: **, p ≤ 0.001: ***, p ≤ 0.0001: ****).

#### Rule-based Finite Element Modeling of Multiscale Intercalated Disc Structure

These ultrastructural TEM data were used to generate populations of rule-based finite element models mimicking ID ultrastructure, incorporating structural features from micro to nano scales (Figure 4A-B). Utilization of specific ultrastructural measurements are illustrated on representative atrial and ventricular ID finite element meshes (Figure 4A-B, Supplementary Table 1). At the largest scale, ventricular cardiomyocytes were longer, wider, deeper, and had greater volume compared to atrial cardiomyocytes. Thus, ventricular IDs had proportionally greater diameter and volume compared to atrial IDs (Figure 4C). Focusing on plicate subdomains, structural features (fraction of ID comprised of plicate subdomains, plicate subdomain length, fraction of plicate subdomains comprised by mechanical junctions, and plicate membrane fold frequency) were largely similar between the two chambers. However, plicate subdomains in ventricles displayed larger membrane fold amplitude, mechanical junction length, and intermembrane spacing at/near mechanical junctions when compared to atria (Figure 4D). Turning to interplicate subdomains, these regions had similar interplicate domain length and comprised similar fractions of IDs in both chambers. However, gap junctions were larger and comprised a greater fraction of the interplicate subdomains in the ventricles relative to atria (Figure 4E). Likewise, intermembrane spacing outside interplicate gap junctions was also greater in the ventricles. These data suggest multiscale differences in ID ultrastructure that may underlie conduction differences between atrial and ventricular myocardium.

**Figure 4.**
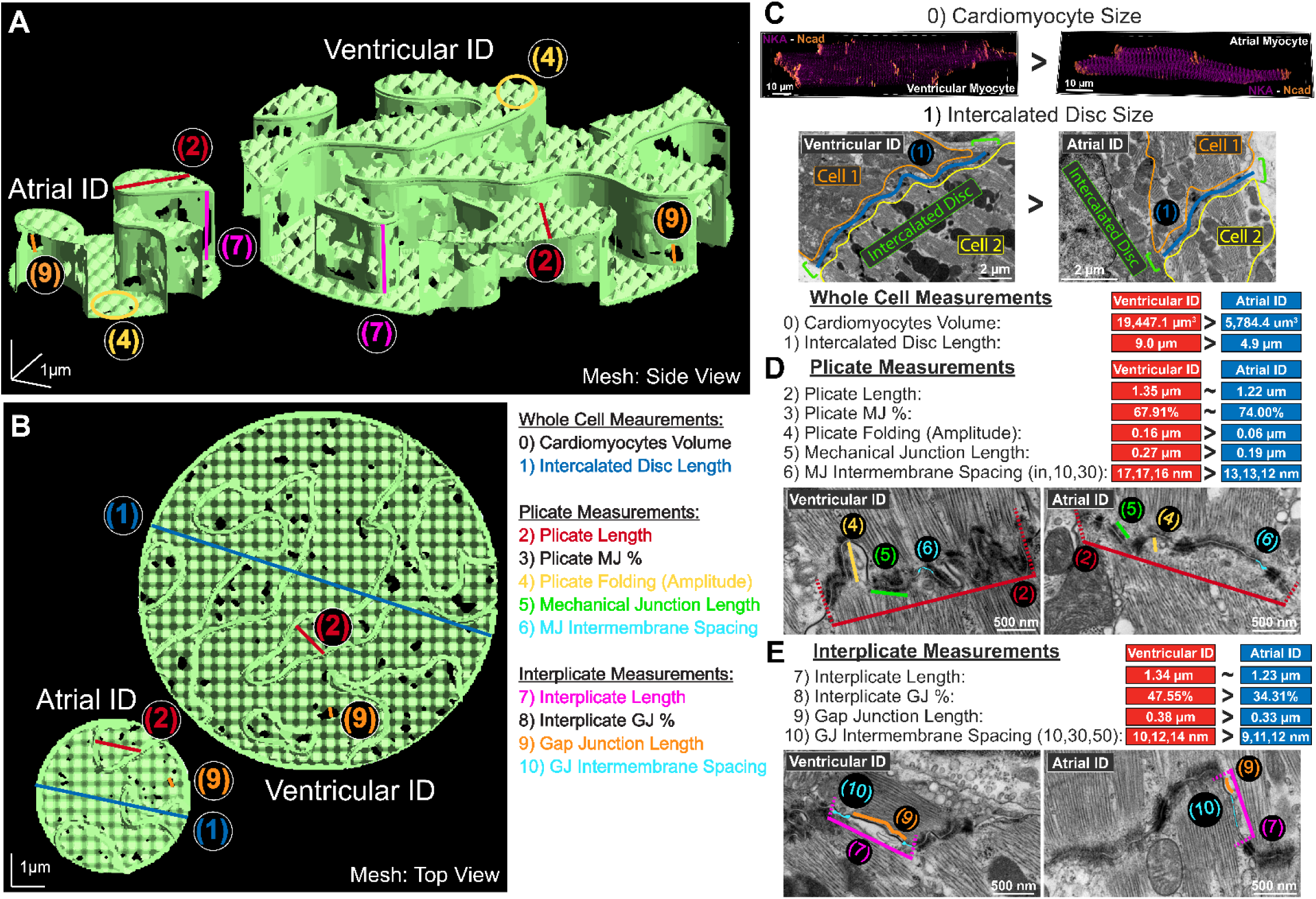
Integration of intercalated disc ultrastructure into rule-based finite element model. Representative finite element meshes of atrial and ventricular intercalated discs: **A)** side view and **B)** top view. Overlay of multiscale intercalated disc measurements used for mesh generation: 0) cardiomyocyte volume, 1) intercalated disc size (blue), 2) plicate length (red), 3) plicate MJ%, 4) plicate folding (amplitude, yellow), 5) mechanical junction length (green), 6) MJ intermembrane spacing (teal), 7) interplicate length (pink), 8) interplicate GJ%, 9) gap junction length (orange), 10) GJ intermembrane spacing (teal). Ultrastructural measurements are annotated in TEM images and summarized as comparison of medians for **C)** whole cell, **D)** plicate, and **E)** interplicate measurements.

### Cellular Scale Distribution of Electrogenic Proteins

The relative distribution of electrogenic proteins (Na_V_1.5 [green], K_ir_2.1 [pink], and NKA [purple]) between the ID and other myocyte regions was assessed by confocal microscopy in isolated atrial and ventricular myocytes with the mechanical junction protein, N-cadherin (N-cad; orange), serving as a marker for the ID (Supplementary Figure 5). Greater abundance of immunosignals for Na_V_1.5, K_ir_2.1, and NKA was observed in atrial myocytes compared to ventricular cells (Supplementary Figure 5G-I; n= 50[A], 50[V] cells for each electrogenic protein from 4 hearts). Na_V_1.5 was enriched at the ID to similar degrees in both atrial and ventricular cardiomyocytes. In contrast, atrial cardiomyocytes had greater ID enrichment of K_ir_2.1 and NKA compared to ventricular cardiomyocytes.

### Spatial Distribution of Junctional and Electrogenic Proteins within the Intercalated Disc

Spatial organization of ID mechanical and electrical junction proteins was further characterized using stochastic optical reconstruction microscopy (STORM). First, we examined the distribution of junctional proteins relative to each other. Consistent with our TEM results, mechanical junction proteins N-cad and Dsp localized to ID regions largely distinct from those containing the gap junction protein, Cx43 (Supplementary Figure 6; n= 5[A], 5[V] images per heart from 3 hearts). Overall, Cx43 was further separated from Dsp (Supplementary Figure 6E, H, Supplementary Figure 7) but located closer to N-cad (Supplementary Figure 6D, G, Supplementary Figure 7) within atrial IDs, compared to ventricular, while the relative distribution of N-cad and Dsp was similar between chambers (Supplementary Figure 6F, I, Supplementary Figure 7).

Next, we assessed the distribution of electrogenic proteins Na_V_1.5, K_ir_2.1, and NKA relative to junctional proteins, Cx43, N-cad, and Dsp. By visual inspection, Na_V_1.5 was identified to co-distribute with N-cad and Dsp in the plicate regions (Figure 5B-C, Supplementary Figure 8A-B), while remaining on the periphery of Cx43 clusters (consistent with gap junctions) in the interplicate regions (Figure 5A, Supplementary Figure 8C). Atrial IDs displayed closer association of Na_V_1.5 with all three junctional proteins compared to ventricular IDs, the difference being most pronounced near N-cad (Figure 5J-L, Supplementary Figure 8D-I; n= 5[A], 5[V] images per heart from 3 hearts). Similar to Na_V_1.5, K_ir_2.1 co-distributed with N-cad and Dsp in the plicate regions (Figure 5E-F, Supplementary Figure 9A-B) but remained on the periphery of Cx43 clusters (Figure 5D, Supplementary Figure 9C). K_ir_2.1 clusters associated largely similarly with all three junctional molecules in both atrial and ventricular IDs with only minor differences observed (Figure 5J-L, Supplementary Figure 9D-I; n= 5[A], 5[V] images per heart from 3 hearts). Lastly, NKA also co-distributed with N-cad and Dsp, while remaining at the periphery of Cx43 clusters (Figure 5G-I, Supplementary Figure 10A-C). NKA associated more closely with N-cad and Cx43 and less closely with Dsp in atrial IDs compared to ventricular (Figure 5J-L, Supplementary Figure 10D-I; n= 5[A], 5[V] images per heart from 3 hearts). In addition, Na_V_1.5, K_ir_2.1, and NKA clusters displayed lower density and volume as well as less variability in these parameters within atrial IDs relative to ventricular IDs (Supplementary Figure 11A-C, 12A-C, 13A-C). In both chambers, larger, denser clusters were preferentially localized near junctions.

**Figure 5.**
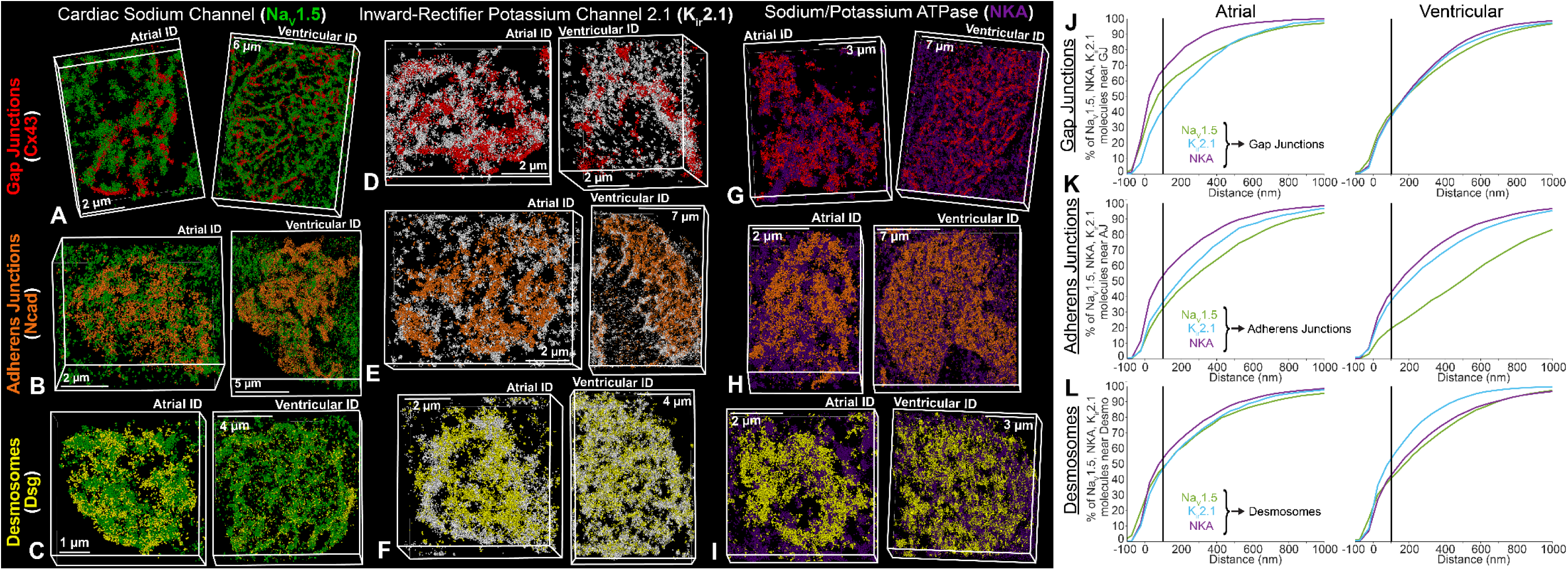
Na_V_1.5, K_ir_2.1, NKA channel distribution relative to junctional proteins. Representative STORM images of atrial and ventricular intercalated disc labeled for protein components of cardiac isoform of the voltage-gated sodium channel (Na_V_1.5; green), inward-rectifier potassium channel 2.1 (K_ir_2.1; white), sodium potassium ATPase (NKA; purple), gap junctions (Cx43; red), adherens junctions (Ncad; orange), and desmosomes (Dsp; yellow). **A)** cardiac isoform of the voltage-gated sodium channel + gap junctions, **B)** cardiac isoform of the voltage-gated sodium channel + adherens junctions, **C)** cardiac isoform of the voltage-gated sodium channel + desmosomes, **D)** inward-rectifier potassium channel 2.1 + gap junctions, **E)** inward-rectifier potassium channel 2.1 + adherens junctions, **F)** inward-rectifier potassium channel 2.1 + desmosomes, **G)** sodium potassium ATPase + gap junctions, **H)** sodium potassium ATPase + adherens junctions, **I)** sodium potassium ATPase + desmosomes. Images were presented as whole intercalated discs. Cumulative distributions of electrogenic proteins (Na_V_1.5 [green], K_ir_2.1 [light blue], and NKA [purple]) relative to intercalated disc junctions: **J)** gap junctions, **K)** adherens junctions, and **L)** desmosomes. Summary plots show molecule-wise cumulative distributions (n= 5[A], 5[V] images/heart from 3 hearts).

Lastly, we assessed the localization of Na_V_1.5, K_ir_2.1, and NKA relative to each other. Na_V_1.5 clusters were mixed with clusters of both NKA and K_ir_2.1 with surrounding clusters predominantly composed of NKA and K_ir_2.1 (Figure 6A-B, Supplementary Figure 14A-B). Similarly, NKA and K_ir_2.1 organized into mixed clusters but with NKA extending more diffusely in space than K_ir_2.1 (Figure 6C, Supplementary Figure 14C). Na_V_1.5 was more closely associated to NKA and K_ir_2.1 in atrial IDs than ventricular (Figure 6D, Supplementary Figure 14D-E, G-H; n= 5[A], 5[V] images per heart from 3 hearts). Atrial and ventricular IDs displayed similar distances between K_ir_2.1 and NKA (Figure 6E, Supplementary Figure 14F, I). Na_V_1.5, K_ir_2.1, and NKA clusters displayed lower density and volume as well as less variability in these parameters within atrial IDs relative to ventricular IDs (Supplementary Figure 15A-C). In both chambers, larger, denser clusters of electrogenic proteins were preferentially localized near each other.

**Figure 6.**
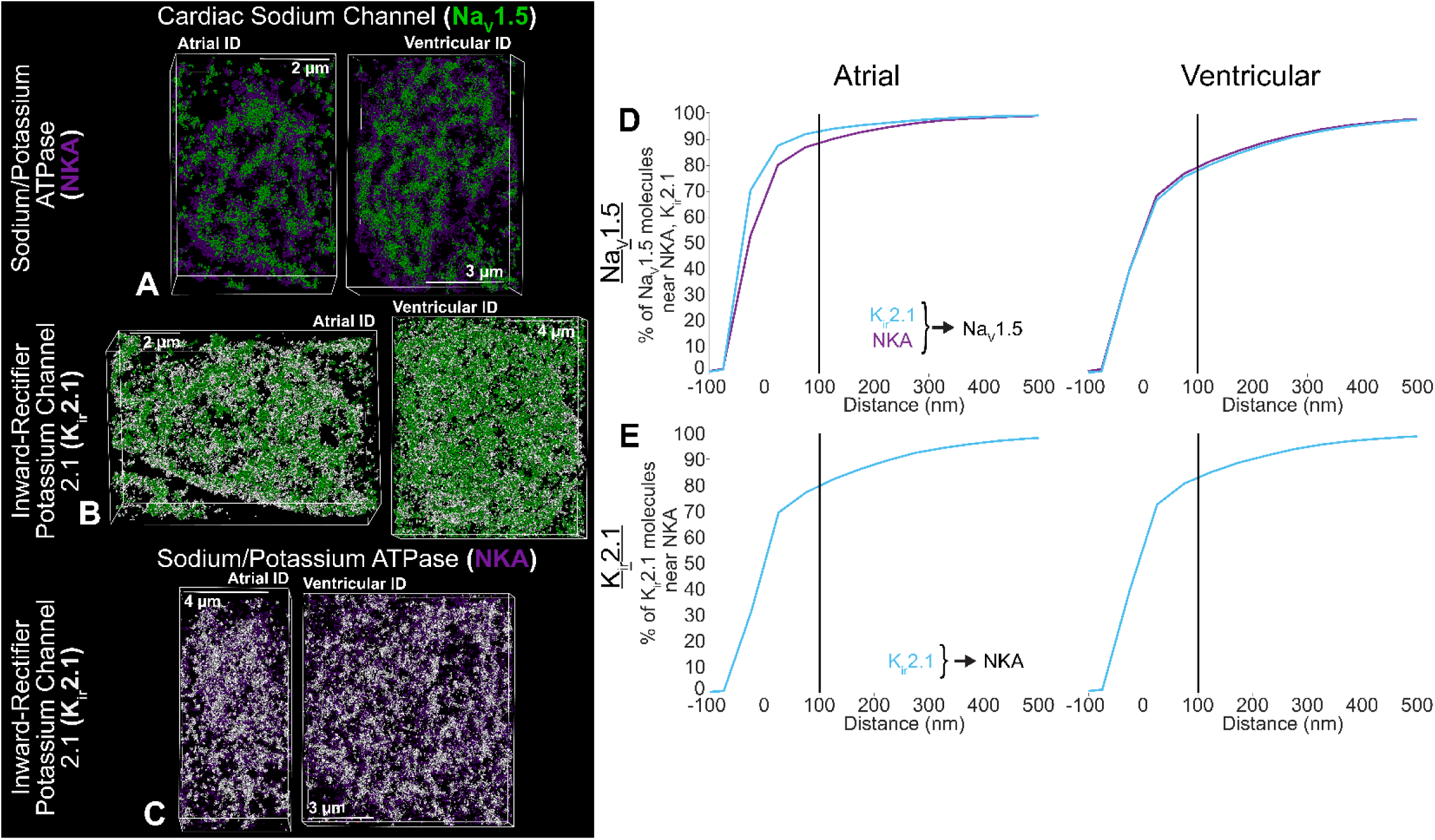
Distribution of electrogenic proteins relative to each other. Representative STORM images of atrial and ventricular intercalated disc labeled for protein components of cardiac isoform of the voltage-gated sodium channel (Na_V_1.5; green), inward-rectifier potassium channel (K_ir_2.1; white), and sodium potassium ATPase (NKA; purple). **A)** cardiac isoform of the voltage-gated sodium channel + sodium potassium ATPase, **B)** cardiac isoform of the voltage-gated sodium channel + inward-rectifier potassium channel, and **C)** sodium potassium ATPase + inward-rectifier potassium channel. Cumulative distributions of electrogenic proteins (K_ir_2.1 [light blue], and NKA [purple]) relative to each other: **D)** Na_V_1.5 relative to K_ir_2.1 and NKA, **E)** K_ir_2.1 relative to NKA. Summary plots show molecule-wise cumulative distributions (n= 5[A], 5[V] images/heart from 3 hearts).

#### Rule-based Finite Element Modeling of Intercalated Disc Molecular Organization

Spatial distributions of electrogenic proteins (Na_V_1.5, K_ir_2.1, NKA) relative to ID junctional landmarks were next incorporated into our rule-based finite element models of atrial and ventricular IDs (Figure 7A-C, Supplementary Table 2). The ultrastructural landmarks (mechanical and electrical junctions) identified by TEM served as reference points to distribute the electrogenic proteins throughout the finite element meshes (Figure 7B). Overall organization of electrogenic proteins relative to junctional landmarks (Figure 5J-L, 7D) and each other (Figure 6D-E, 7E) was captured in the form of probability density functions (Figure 7D-E) and cumulative distributions (Figure 5J-L, 6D-E) of distances within the ID from both chambers.

**Figure 7.**
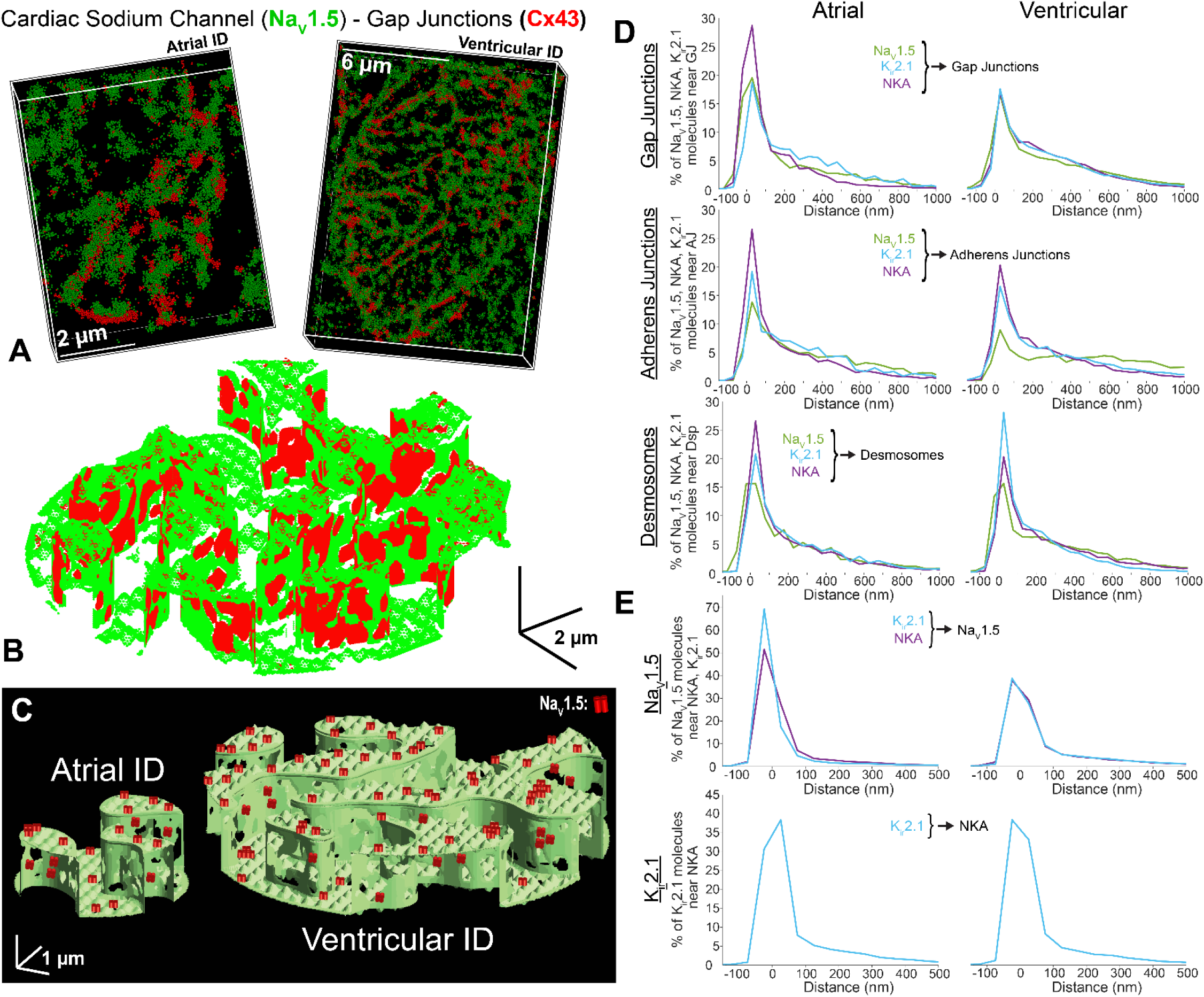
Integration of intercalated disc protein organization into rule-based finite element model. Pipeline for integrating protein organization: **A)** STORM-based assessment of electrogenic protein distribution relative to junctional landmarks, **B)** *in silico* mapping of electrogenic protein placement relative to landmarks previously established in ultrastructural meshes via TEM, **C)** cartoon of electrogenic protein integration into atrial (left) and ventricular (right) ultrastructural meshes. Summary plots of distribution of electrogenic proteins (Na_V_1.5 [green], K_ir_2.1 [light blue], and NKA [purple]) **D)** relative to intercalated disc junctions (gap junctions, adherens junctions, and desmosomes) and **E)** relative to each other. Summary plots show molecule-wise probability distribution functions.

Electrogenic proteins were more closely associated with gap junctions and adherens junctions within atrial IDs, when compared to ventricular IDs (Figure 5J-L, 7D). Atrial IDs also displayed greater enrichment of electrogenic proteins near gap junctions with NKA being closest, followed by Na_V_1.5 and K_ir_2.1 (Figure 5J). In contrast, the electrogenic proteins displayed similar patterns of localization relative to adherens junctions and desmosomes in both chambers (Figure 5K-L, 7D). NKA was closest to adherens junctions, followed by K_ir_2.1, and Na_V_1.5, while all three were similarly distributed relative to desmosomes. Na_V_1.5 channels were more closely associated with NKA and K_ir_2.1 within atrial IDs, when compared to ventricular. Atrial IDs displayed higher prevalence of Na_V_1.5 closer to K_ir_2.1 followed by NKA, while ventricular IDs maintained similar Na_V_1.5 distributions relative to NKA and K_ir_2.1 (Figure 6D, 7E). There were no discernable differences between atrial and ventricle IDs with respect to the association of K_ir_2.1 to NKA (Figure 6E, 7E).

### Functional Implications of Intercalated Disc Structural Organization

To experimentally validate structure-function relationships predicted by our model, we performed whole-heart optical mapping experiments in Langendorff-perfused mouse hearts. Isochrone maps of activation (Figure 8A-B) and action potential traces (Figure 8C) illustrate faster conduction in the ventricles than the atria. Consistent with previous studies^32^, both longitudinal and transverse conduction velocities were greater in the ventricles in comparison to the left atrium (Figure 8D-E; Supplementary Table 3, from 3 hearts). Simulations were performed of both atrial and ventricular tissue, integrating measures of cell geometry, ID nanoscale structure, and electrogenic protein organization. Importantly, experimentally observed longitudinal conduction velocities (Figure 8D) displayed close agreement with the computationally predicted values (Figure 8F-G), providing direct validation of the model.

**Figure 8.**
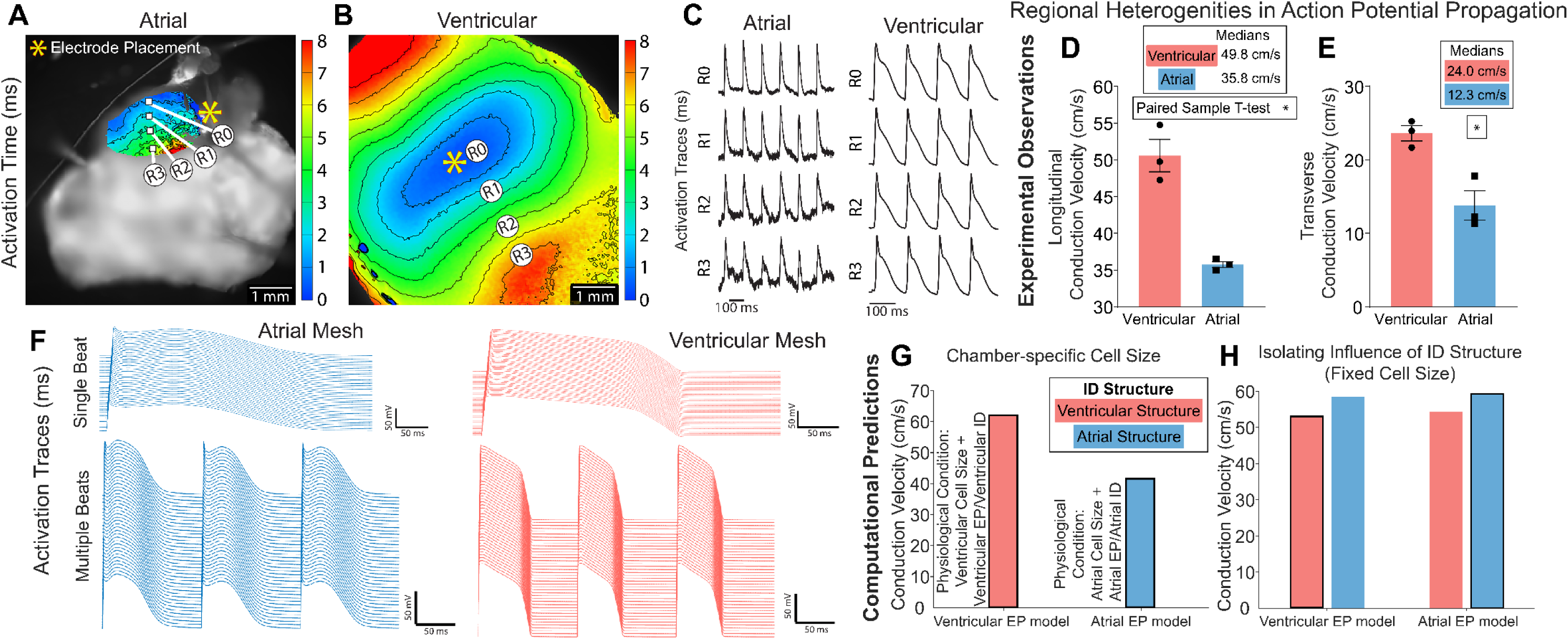
Functional implications of intercalated disc structural heterogeneities. Representative isochrone maps of activation from optical mapping of **A)** atrial and **B)** ventricular myocardium with **C)** action potential duration (APD) traces matched to sites overlaid on the image. Propagation across the tissue was captured as **D)** longitudinal conduction velocity and **E)** transverse conduction velocity (from 3 hearts; p ≤ 0.5: *, paired-sample T-test). Functional effects of incorporating atrial and ventricular cellular and intercalated disc structure into the rule-based finite element model **F-H)**. Computational model generated **F)** action potential traces and longitudinal conduction velocity with **G)** chamber-specific cell size and **H)** uniform cell length of 100 μm. Outlined bars represent chamber-specific structures integrated with corresponding EP models (i.e. Atrial structure with atrial EP model).

Finally, we sought to delineate the specific roles of inter-chamber differences in macro/cellular scale structure vs. ID micro/nano structure and molecular organization in determining conduction differences between atria and ventricle. We highlight that this is not experimentally feasible, as myocyte geometry, ID structure, and ionic current properties all vary concurrently between the two chambers. To undertake this investigation, we performed tissue simulations with fixed myocyte geometry (i.e., equal between atria and ventricles), while retaining other differences in ID structure and simulated tissues with each ID structure and with either atrial or ventricular electrophysiology models. In these simulations, we observed that atrial conduction velocity was faster, independent of the electrophysiology model governing lateral and ID membrane ionic currents (Figure 8H; Supplementary Table 3). Thus, even though on a tissue level atrial conduction is slower compared to ventricular tissue due to the effects of myocyte geometry, atrial IDs are able to support faster conduction than ventricular IDs.

## DISCUSSION

Structural differences between atrial and ventricular myocardium are well established at the organ (eg. wall thickness), tissue, and cellular levels (eg. cell size^18–22^) and are linked to differences in action potential propagation under normal physiological conditions, as well as arrhythmia susceptibility under pathological conditions. However, there is limited data available on structural differences at subcellular scales, particularly vis-à-vis ultrastructure and protein organization at sub-micron scales. Previous studies have linked disruption of ID nanodomains with severe functional consequences over both acute (de-adhesion, edema^4, 8^) and chronic (arrhythmogenic cardiomyopathy, heart failure, atrial fibrillation^33–38^) time scales. As a key first step towards understanding how ID structure determines function in health and disease, we undertook a systematic comparative study of ID properties in healthy murine atria and ventricles with an eye towards elucidating differences in action potential propagation. By combining experimental and computational modeling approaches, we demonstrate that, once inter-chamber differences in myocyte geometry are accounted for, atrial IDs possess ultrastructural and molecular organization properties conducive to faster action potential propagation in comparison to their ventricular counterparts.

### Intercalated Disc Ultrastructure

Although it has been known for some time that atrial and ventricular IDs differ at the ultrastructural level, previous studies have evaluated specific aspects in isolation and mostly focused on either chamber without direct comparisons between atrial and ventricular IDs. Thus, we systematically compared atrial and ventricular ID ultrastructure using electron microscopy. However, the functional implications of such differences cannot be considered without also considering differences in cardiomyocyte geometry, a key determinant of conduction velocity. Consistent with previous studies in rabbits^39^ and humans^27^, we found murine atrial cardiomyocytes to be smaller in length, width, depth, and volume than ventricular cardiomyocytes. Next, we systematically assessed ID structural properties ranging from overall size at the microscale down to intermembrane spacing at the nanoscale (Supplementary Table 1) and compared our results with previous reports, where available. Overall, atrial IDs were smaller than ventricular IDs; however, the fractions of IDs comprised by plicate and interplicate regions were similar between chambers. Atrial IDs had smaller mechanical junctions and tighter intermembrane spaces near both mechanical and gap junctions compared to ventricular IDs. Within plicate subdomains, we found no significant difference between chambers in membrane fold frequency, as previously reported in humans^27^. We also measured fold amplitudes similar to previous measurements in murine ventricular tissue^40–42^, and mechanical junction lengths and intermembrane distances similar to previous data from human ventricular tissue^35^. Within interplicate subdomains, we measured similar gap junction lengths to previous data from human ventricular tissue^35^. In addition to highlighting chamber-specific differences, the close alignment between our ultrastructural measurements in mice and previous studies from other mammalian species may suggest a degree of evolutionarily conserved ID properties. It should be noted, however, that certain properties, such as the distribution of gap junctions within the ID, show species-specific differences, particularly between rodents and primates^43^.

### Intercalated Disc Molecular Organization

Previous studies indicate that cardiac conduction is strongly influenced by ion channel localization to specialized ID nanodomains^3–7^. Therefore, we characterized the molecular organization of key electrogenic proteins directly associated with cardiac excitability, Na_V_1.5, K_ir_2.1, and NKA, at the cellular and ID levels. Consistent with previous studies in multiple mammalian species, we observed Na_V_1.5^6, 44–47^, K_ir_2.1^6, 48–53^, and NKA (α1 subunit)^11, 54–56^ localizing to ID, lateral sarcolemma and transverse tubules using confocal microscopy, with Na_V_1.5, in particular, showing marked enrichment at the ID. However, these previous reports were largely qualitative, precluding direct comparisons with our quantitative measurements. Interestingly, we observed more abundant immunosignals for Na_V_1.5, K_ir_2.1, and NKA at both ID and non-ID sites in atrial cardiomyocytes than in ventricular cardiomyocytes.

Previous studies have established the local enrichment of Na_V_1.5 near adherens junctions and gap junctions in ventricular tissue / cardiomyocytes from mice, guinea pigs, and rats^4, 24, 44, 57, 58^. Furthermore, nanoscale shifts in ion channel localization within the ID and/or disruption of ion channel-rich nanodomains have been linked to proarrhythmic conduction defects in both acute^8^ and chronic disease states^33, 34, 36^. Yet, little is known about the intra-ID distributions of K_ir_2.1 and NKA and their spatial relationship to Na_V_1.5. Therefore, we used super-resolution microscopy to systematically assess the localization of these electrogenic proteins to gap junction and mechanical junction-associated ID nanodomains. Na_V_1.5 had closer association to gap junctions and mechanical junctions within atrial IDs compared to ventricular. K_ir_2.1 was equally enriched near gap junctions and mechanical junctions in both chambers. NKA associated more closely with adherens junctions and gap junctions within atrial IDs, while in ventricular IDs, NKA associated more closely with desmosomes. Next, we examined the localization of the electrogenic proteins relative to each other: Na_V_1.5 associated more closely with NKA and K_ir_2.1 in atrial IDs than in ventricular, while the localization of K_ir_2.1 relative to NKA was consistent between gap junctions and mechanical junctions in both chambers.

### Functional Implications of Intercalated Disc Structure

It has long been recognized that conduction in the healthy heart is faster in ventricular myocardium than in atrial^32, 59–61, 62, 63^, an observation we recapitulate here. However, these conduction differences have been largely ascribed to differences in myocyte geometry^2^ with little known about the functional implications of inter-chamber differences in ID structure. This stems from current experimental methods lacking the spatial resolution to assess conduction at the scale of individual IDs. While conduction dependence on subcellular structure at such scales has been studied via computational models, most included no representation of the ID and even recent modeling studies have grossly oversimplified ID structure^3, 5, 7, 44, 64, 65^. This was largely a result of available data on multiscale ID structure being sparse, fragmentary and limited to simple measures of central tendency (thus, not capturing complex variability). We provide here the first ever comprehensive quantification of ID structure and molecular organization across multiple spatial scales. Further, we provide measurements from both left atria and left ventricular free wall tissue, laying the groundwork for future efforts to more closely sample spatial heterogeneities.

We also demonstrate here a novel approach to generate populations of IDs *in silico* using quantitative multiscale data from light and electron microscopy and a method to uncover key structure-function relationships by incorporating them into physiological models of propagation. This builds on our recently publication which demonstrated the first computational model to incorporate 3D ID ultrastructure derived from electron microscopy^9^. Even lacking molecular organization data included here, that model revealed previously unappreciated dependence of conduction on ID structure: the presence of larger gap junctions at the ID periphery increased conduction velocity and intermembrane distance in the interplicate regions had a gap junction coupling-dependent biphasic influence on conduction. Here, we present a major step forward, incorporating experimentally-measured cell size, ultrastructure, and protein organization of atrial and ventricular IDs to identify chamber-specific mechanisms of conduction.

These chamber-specific models closely recapitulated optical mapping measurements of atrial and ventricular conduction velocities made under physiological conditions. As an early foray to elucidate conduction dependence on ID structure, we equalized cell size between the ventricular and atrial models. This approach revealed that atrial IDs support faster conduction than ventricular IDs, which is masked by cell size under normal physiological conditions. Identifying important directions for more in-depth mechanistic modeling studies, these studies point to specific aspects of ID structure at different scales that contribute to faster conduction through atrial IDs: Compared to ventricular IDs, atrial IDs maintain tighter intermembrane spaces near mechanical junctions and gap junctions, display greater abundance of Na_V_1.5, K_ir_2.1, and NKA, and have Na_V_1.5 more closely associating with gap junctions and mechanical junctions. While increased Na_V_1.5 expression is an established contributor to faster conduction, previous studies from us and others^3, 5, 7, 66^ have also identified close membrane apposition and Na_V_1.5 enrichment at juxtajunctional ID sites as factors that promote faster conduction. Importantly, disruption of these juxtajunctional ID nanodomains and loss of Na_V_1.5 from such sites have been linked with proarrhythmic conduction defects in both atria and ventricles in animal models as well as human patients^4, 8, 33^. In addition to the aforementioned factors, atrial IDs also have NKA localized closer to adherens junctions and gap junctions, and Na_V_1.5 associated more closely with NKA and K_ir_2.1. Taken together, the ultrastructure and molecular organization of atrial IDs may support more robust excitability, especially at faster heart rates.

Thus, our work provides the first quantitative multiscale dataset capturing ID structure and molecular organization across spatial scales, and demonstrates a pipeline for generating such data and integrating them into mechanistic models to query functional implications. Our early mechanistic studies here highlight previously unrecognized aspects of ID structure, which when remodeled in disease states, can contribute to proarrhythmic conduction defects, motivating more in-depth studies into such structure-function relationships under different conditions of health and disease.

#### Limitations

Given that the present work only utilized male mice, future work is needed to provide detailed comparisons of ID structure between male and female hearts, as well as comparisons with non-murine mammalian species, especially human. Secondly, future work must also expand to include non-myocyte cells, which play key roles modulating the behavior of cardiac myocytes. Our ability to precisely resolve protein localization is limited by linkage error resulting from the use of antibodies. Future work with fluorescently tagged nanobodies can help minimize this issue. Lastly, given that our electron microscopy was performed on thin sections, some variation in measurements such as intermembrane spacing reflects variations in the orientation of ID structures relative to the section plane. However, for the purposes of inter-chamber comparisons, this issue is mitigated by our use of large numbers of measurements from multiple samples. Even so, future work using high resolution volume electron microscopy can directly address this issue, provided that approaches for high throughput segmentation of three-dimensional electron microscopy data are developed.

## CONCLUSIONS

We provide here the first ever systematic quantification of ID ultrastructure and molecular organization from healthy adult murine atria and ventricles. Our combined experimental and computational modeling studies revealed that atrial IDs can support faster conduction than ventricular IDs, which is normally masked by inter-chamber differences in myocyte geometry. Further, we identify key ultrastructural and molecular organization features underpinning the ability of atrial IDs to support faster conduction. These data serve as the first stepping stone to elucidating chamber-specific impacts of pathological ID remodeling as occurs in many arrhythmic diseases.

## ACKNOWLEDGEMENTS

The authors would like to thank Dr. Sara Cole, Ms. Sarah Mikula, Mr. Jeff Tonniges, and Ms. Kendall Gallagher from the OSU Campus Microscopy & Imaging Facility for preparing the electron microscopy samples for this study, Drs. Sandor Györke, Thomas Hund and Dmitry Terentyev for fruitful discussions. The authors also thank their friends, Fritz and Zingiber, for their valuable support through the conduct of this study.

## SUPPLEMENTARY RESULTS

**Supplementary Figure 1.**
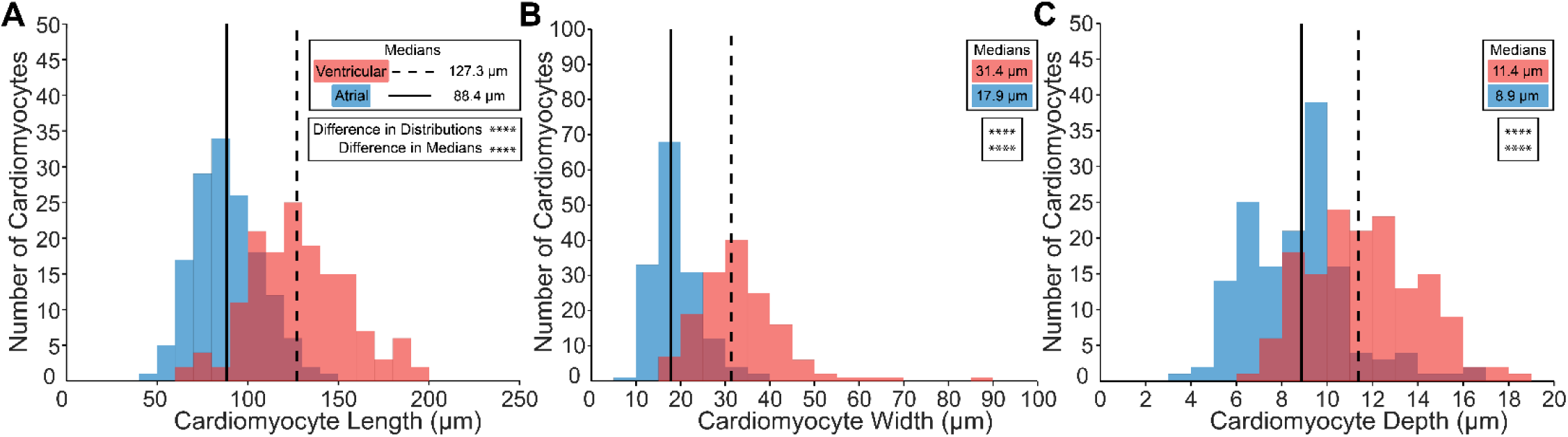
Cardiomyocyte Dimensions. Cardiomyocyte **A)** length, **B)** width, and **C)** depth (n= 151[A], 150[V] measurements from 4 hearts). Distributions were represented as red for ventricular and blue for atrial. Medians were represented as dashed line for ventricular and solid line for atrial. Differences in distributions and medians were statistically proven with two-sample Kolmogorov-Smirnov test and Wilcoxon signed-rank test, respectively (p > 0.05: ns, p ≤ 0.05: *, p ≤ 0.01: **, p ≤ 0.001: ***, p ≤ 0.0001: ****).

**Supplementary Figure 2.**
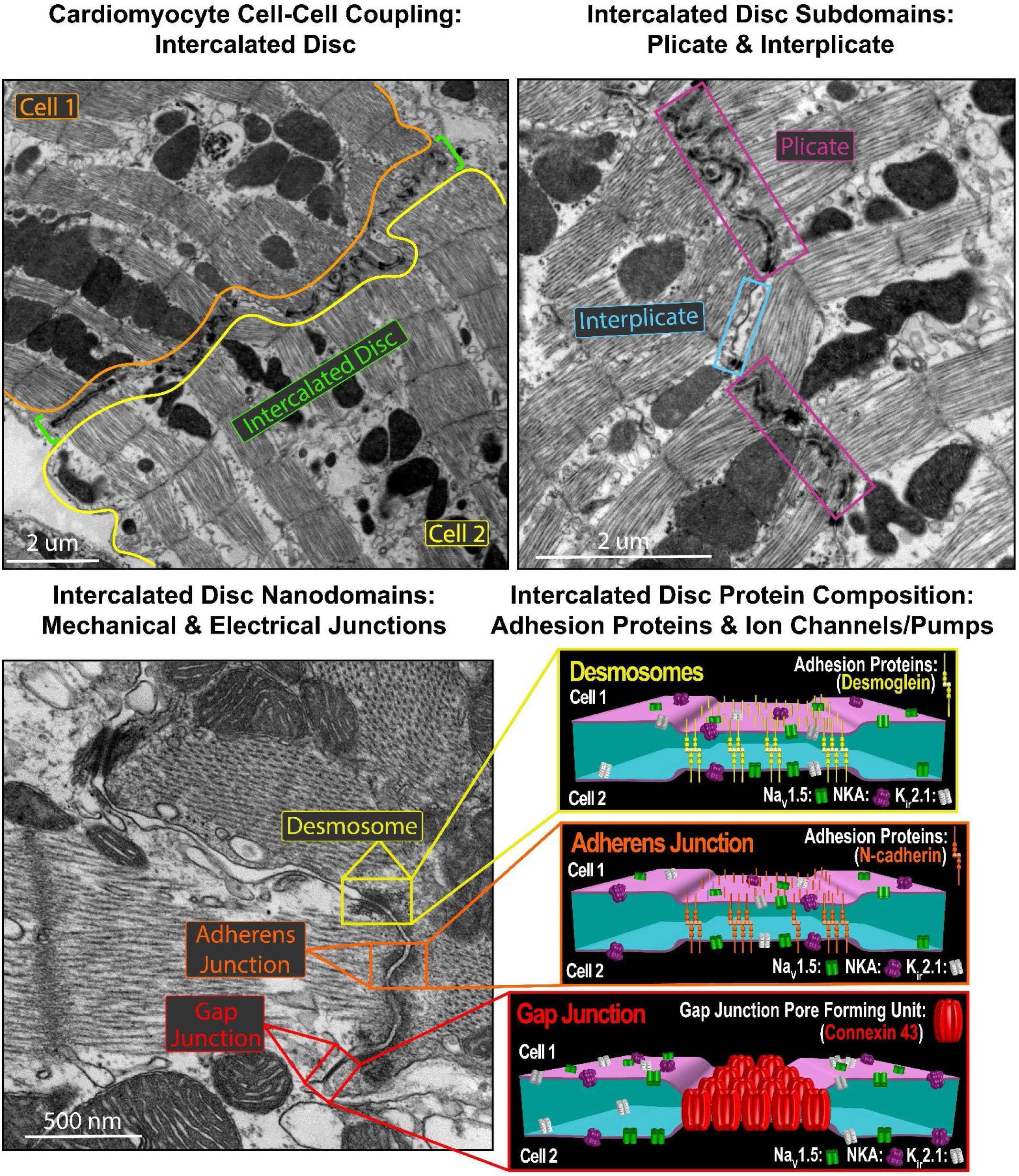
Intercalated Disc Ultrastructure. Representative TEM images annotated to highlight intercalated disc subdomains, nanodomains, and protein composition. Labels are in structure-specific colors: intercalated disc (green brackets), plicate (magenta box), interplicate (light blue box), adherens junction (orange box), desmosome (yellow box), gap junction (red box). Cartoons highlight landmark proteins used to identify intercalated disc junctions (adherens junctions marked by N-cadherin [orange], desmosomes marked by desmoglein [yellow], and gap junctions marked by Connexin 43 [red]) and surrounding electrogenic proteins (cardiac isoform of the voltage-gated sodium channel [Na_V_1.5; green channel] inward-rectifier potassium channel 2.1 [K_ir_2.1; white channel], and sodium potassium ATPase [NKA; purple channel]).

**Supplementary Figure 3.**
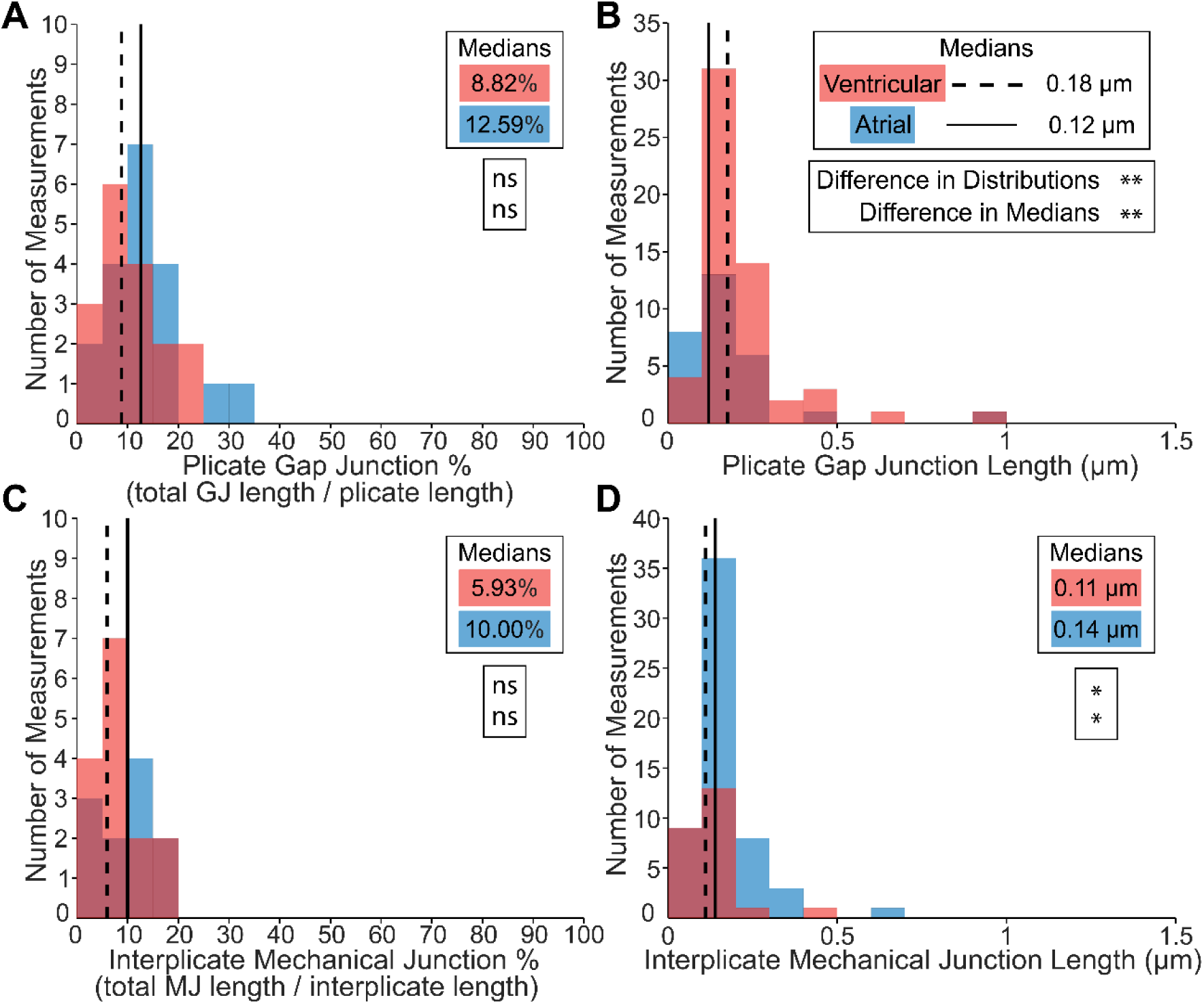
Non-predominant intercalated disc nanodomain structures in the plicate and interplicate subdomains. Plicate gap junctions were characterized by **A)** percentage relative to the plicate region (n= 19[A], 17[V] measurements), and **B)** length (n= 29[A], 56[V] measurements). Interplicate mechanical junctions were characterized by **C)** percentage relative to the interplicate regions (n= 11[A], 15[V] measurements), and **D)** length (n= 57[A], 24[V] measurements). Performed on 3 hearts. Distributions were represented as red for ventricular and blue for atrial. Medians were represented as dashed line for ventricular and solid line for atrial. Differences in distributions and medians were statistically proven with two-sample Kolmogorov-Smirnov test and Wilcoxon signed-rank test, respectively (p > 0.05: ns, p ≤ 0.05: *, p ≤ 0.01: **, p ≤ 0.001: ***, p ≤ 0.0001: ****).

**Supplementary Figure 4.**
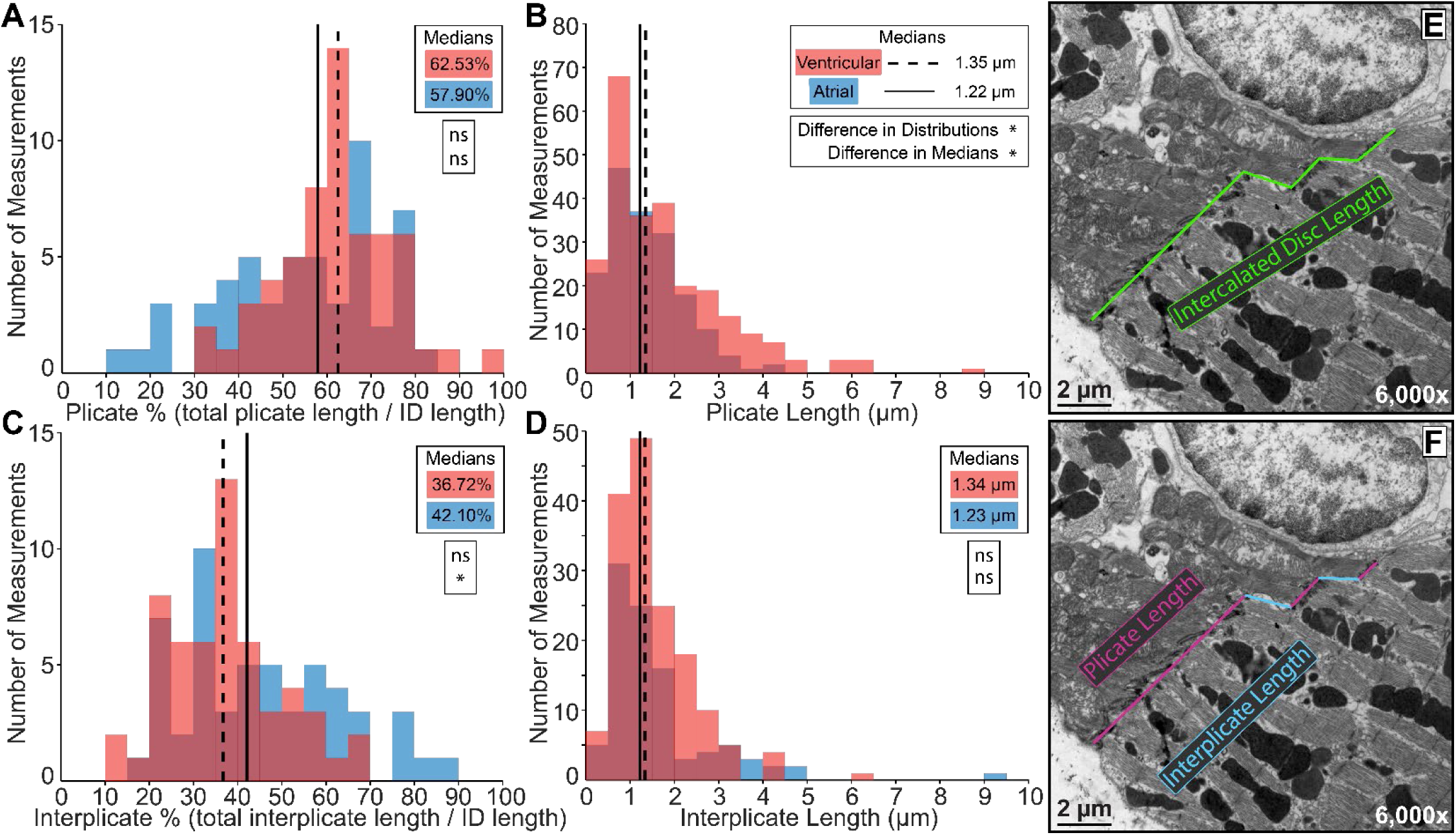
Intercalated Disc Subdomain Structures: Plicate vs Interplicate. Representative TEM images annotated to show intercalated disc subdomain structural measurements: **E)** intercalated disc length (green trace), **F)** plicate length (magenta trace), and interplicate length (light blue trace). Lengths of plicate and interplicate subdomains are shown as **A, C)** percentage relative to the intercalated disc length (plicate n= 53[A], 58[V], and interplicate n= 53[A], 55[V] measurements), and **B, D)** actual length (plicate n= 174[A], 247[V], and interplicate n= 97[A], 160[V] measurements). Performed on 3 hearts. Distributions are represented as red for ventricular and blue for atrial measurements. Medians are represented as dashed line for ventricular and solid line for atrial. Differences in distributions and medians were statistically proven with two-sample Kolmogorov-Smirnov test and Wilcoxon signed-rank test, respectively (p > 0.05: ns, p ≤ 0.05: *, p ≤ 0.01: **, p ≤ 0.001: ***, p ≤ 0.0001: ****).

**Supplementary Figure 5.**
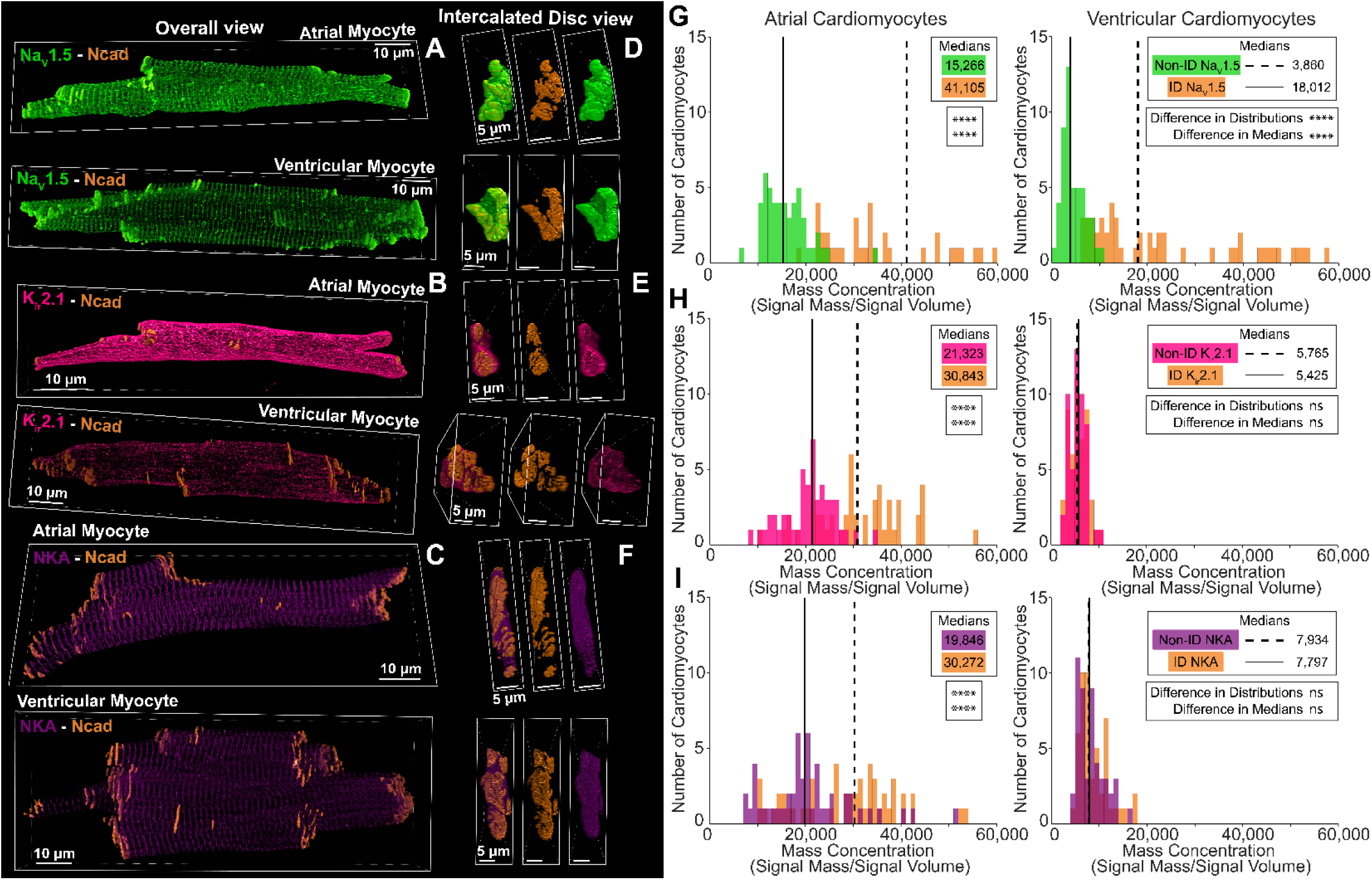
Whole cell distribution of electrogenic protein. Representative 3D confocal images of atrial and ventricular isolated cardiomyocytes stained for intercalated disc marker for adherens junctions (N-cad, orange) and electrogenic proteins: **A, D)** cardiac isoform of the voltage-gated sodium channel (Na_V_1.5; green), **B, E)** inward-rectifier potassium channel 2.1 (K_ir_2.1; pink), and **C, F)** sodium potassium ATPase (NKA; purple). Cardiomyocytes are presented in **A-C)** longitudinal and **D-F)** transverse views. Subcellular distribution of electrogenic proteins was characterized by their **G-I)** relative localization to intercalated disc (orange distributions) and non-intercalated disc regions (green: Na_V_1.5, pink: K_ir_2.1, purple: NKA) (n= 50[A], 50[V] cells for each electrogenic protein). Performed on 4 hearts. Medians were represented as dashed line for ventricular and solid line for atrial. Differences in distributions and medians were statistically proven with two-sample Kolmogorov-Smirnov test and Wilcoxon signed-rank test, respectively (p > 0.05: ns, p ≤ 0.05: *, p ≤ 0.01: **, p ≤ 0.001: ***, p ≤ 0.0001: ****).

**Supplementary Figure 6.**
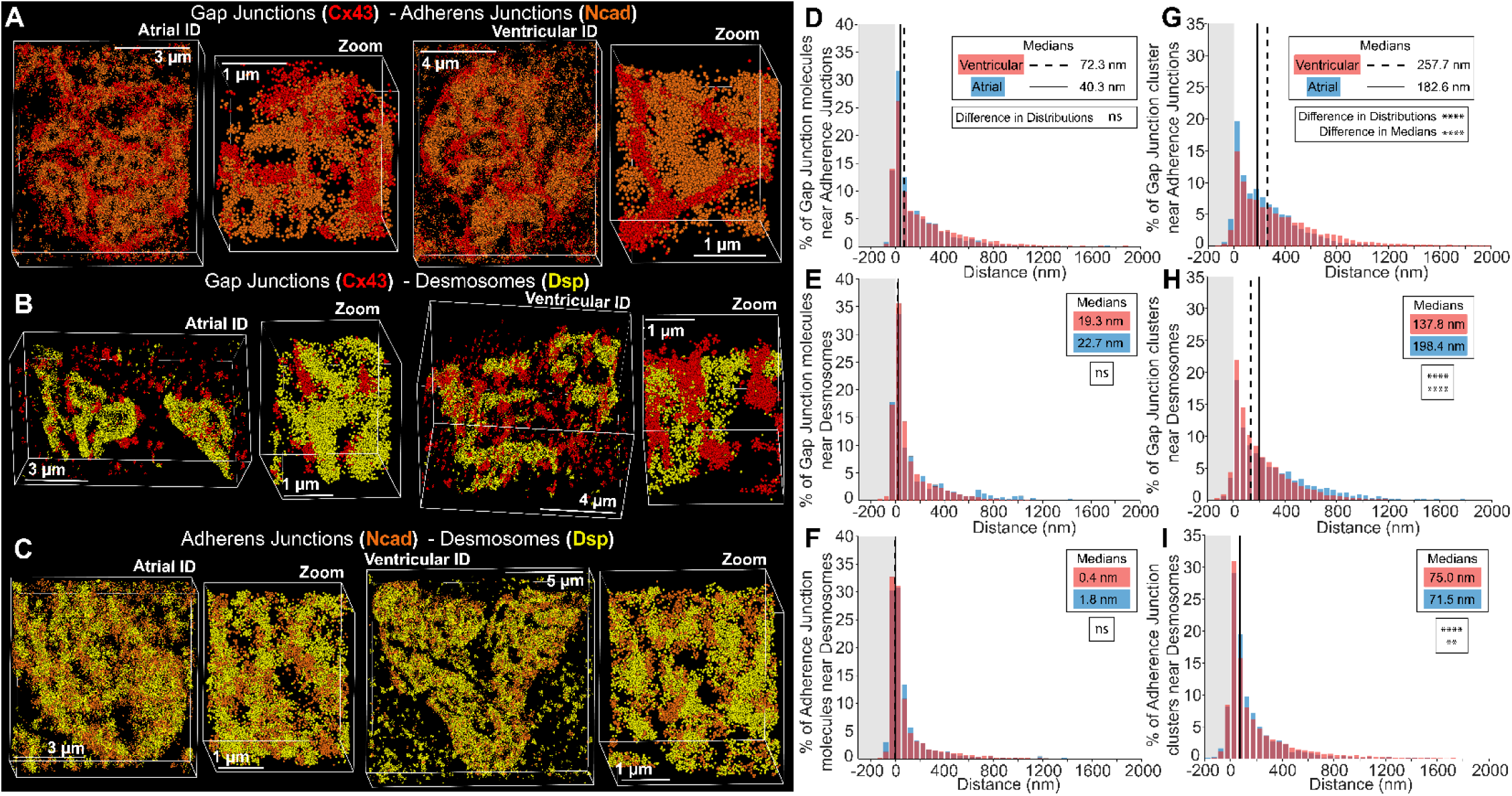
Junctional protein distribution relative to each other. Representative STORM images of atrial and ventricular intercalated disc labeled for proteins components of gap junctions (Cx43; red), adherens junctions (Ncad; orange), and desmosomes (Dsp; yellow). **A)** gap junctions + adherens junctions, **B)** gap junctions + desmosomes, and **C)** adherens junctions + desmosomes. Images were presented as whole intercalated disc and zoomed in views. Protein localization measured as **D-F)** % molecules and **G-I)** % clusters of protein localized relative to the co-labeled protein (n= 5[A], 5[V] images/heart from 3 hearts). Distributions represented in red for ventricular and blue for atrial measurements. Medians were represented as dashed line for ventricular and solid line for atrial. Differences in distributions and medians were statistically proven with two-sample Kolmogorov-Smirnov test and Wilcoxon signed-rank test, respectively (p > 0.05: ns, p ≤ 0.05: *, p ≤ 0.01: **, p ≤ 0.001: ***, p ≤ 0.0001: ****).

**Supplementary Figure 7.**
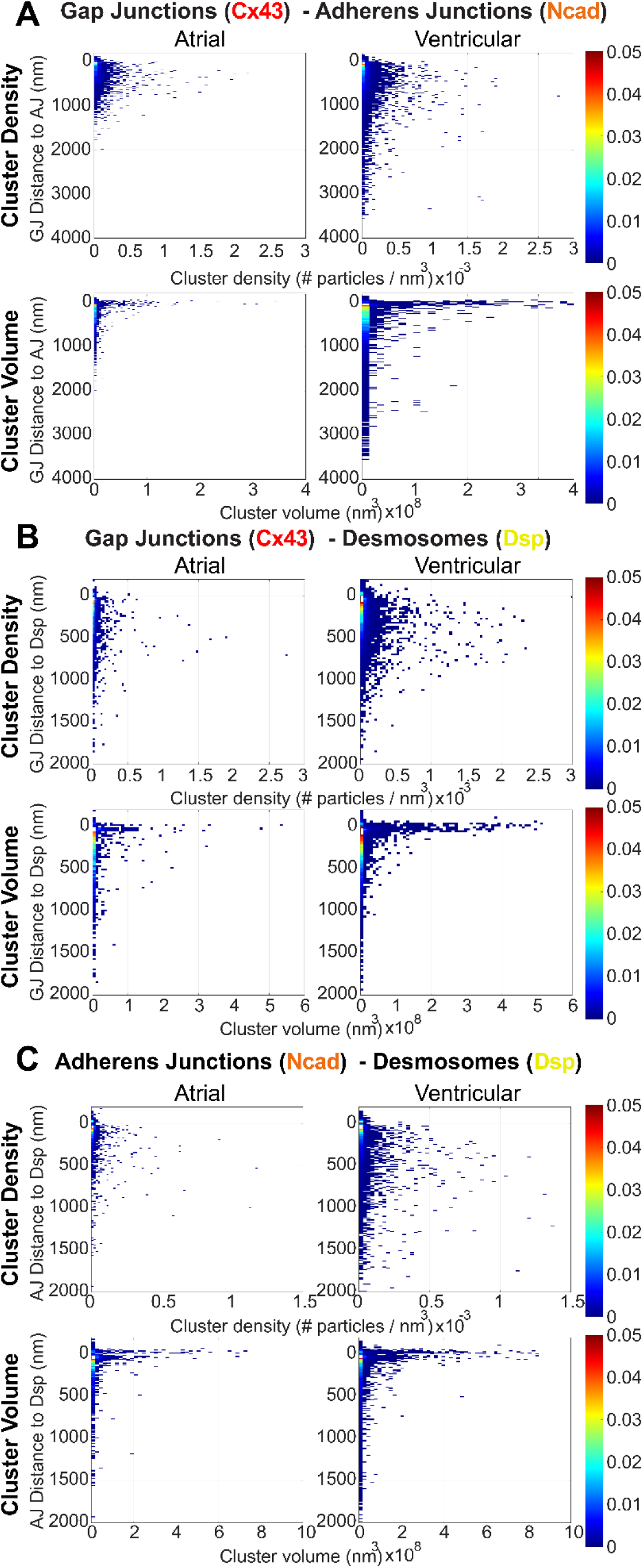
Cluster density and volume for junctional proteins relative to each other. Bivariate histograms of cluster density and volume relative to distance from landmark proteins generated by STORM-based relative localization analysis (STORM-RLA). Analysis of intercalated disc junctional protein clustering: **A)** gap junctions relative to adherens junctions, **B)** gap junctions relative to desmosomes, and **C)** adherens junctions relative to desmosomes.

**Supplementary Figure 8.**
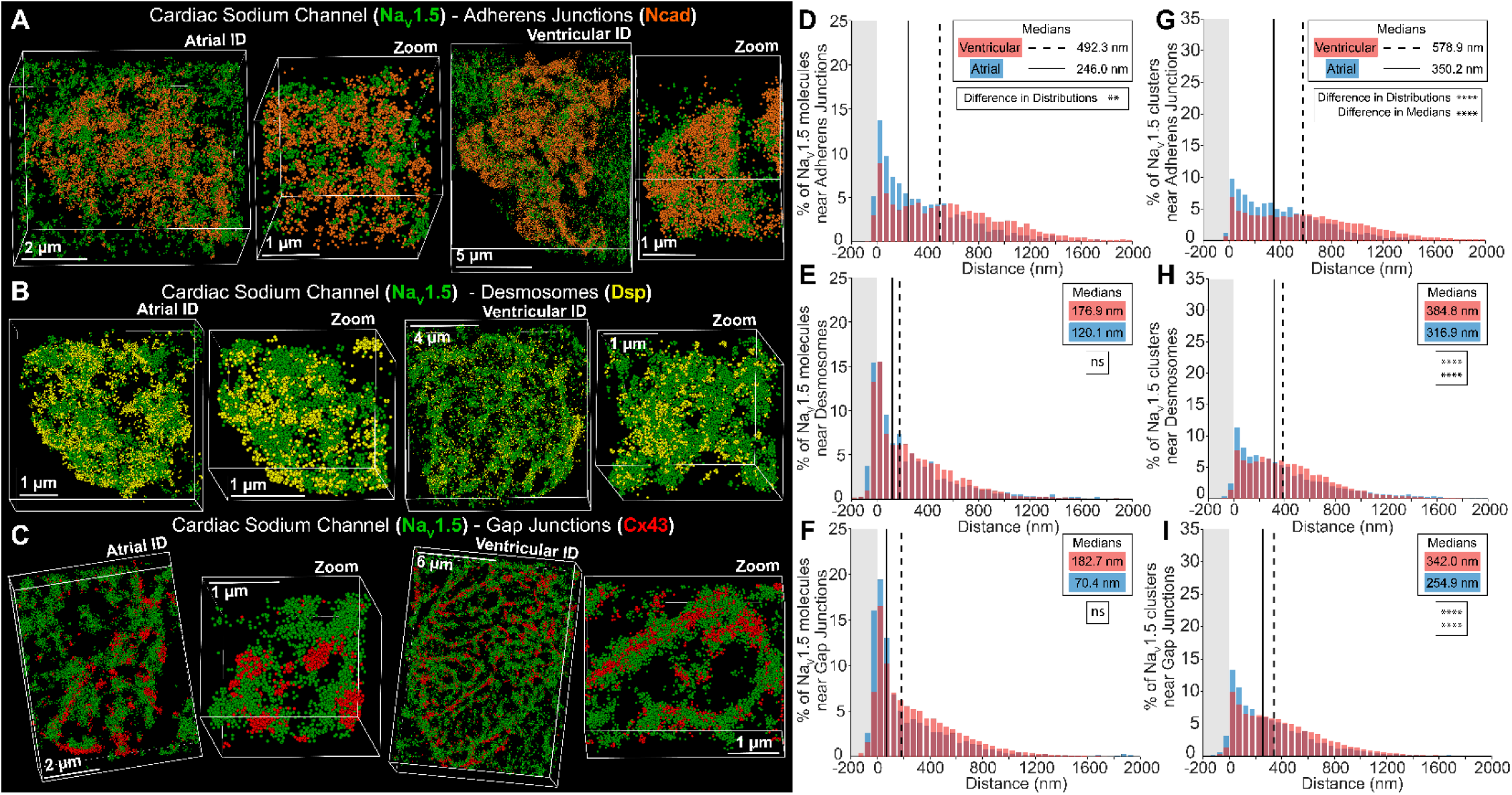
Na_V_1.5 channel distribution relative to junctional proteins. Representative STORM images of atrial and ventricular intercalated disc labeled for protein components of cardiac isoform of the voltage-gated sodium channel (Na_V_1.5; green), gap junctions (Cx43; red), adherens junctions (Ncad; orange), and desmosomes (Dsp; yellow). **A)** cardiac isoform of the voltage-gated sodium channel + adherens junctions, **B)** cardiac isoform of the voltage-gated sodium channel + desmosomes, and **C)** cardiac isoform of the voltage-gated sodium channel + gap junctions. Images were presented as whole intercalated disc and zoomed in views. Protein localization measured as **D-F)** % molecules and **G-I)** % clusters of protein localized relative to the co-labeled protein (n= 5[A], 5[V] images/heart from 3 hearts). Distributions represented in red for ventricular and blue for atrial measurements. Medians were represented as dashed line for ventricular and solid line for atrial. Differences in distributions and medians were statistically proven with two-sample Kolmogorov-Smirnov test and Wilcoxon signed-rank test, respectively (p > 0.05: ns, p ≤ 0.05: *, p ≤ 0.01: **, p ≤ 0.001: ***, p ≤ 0.0001: ****).

**Supplementary Figure 9.**
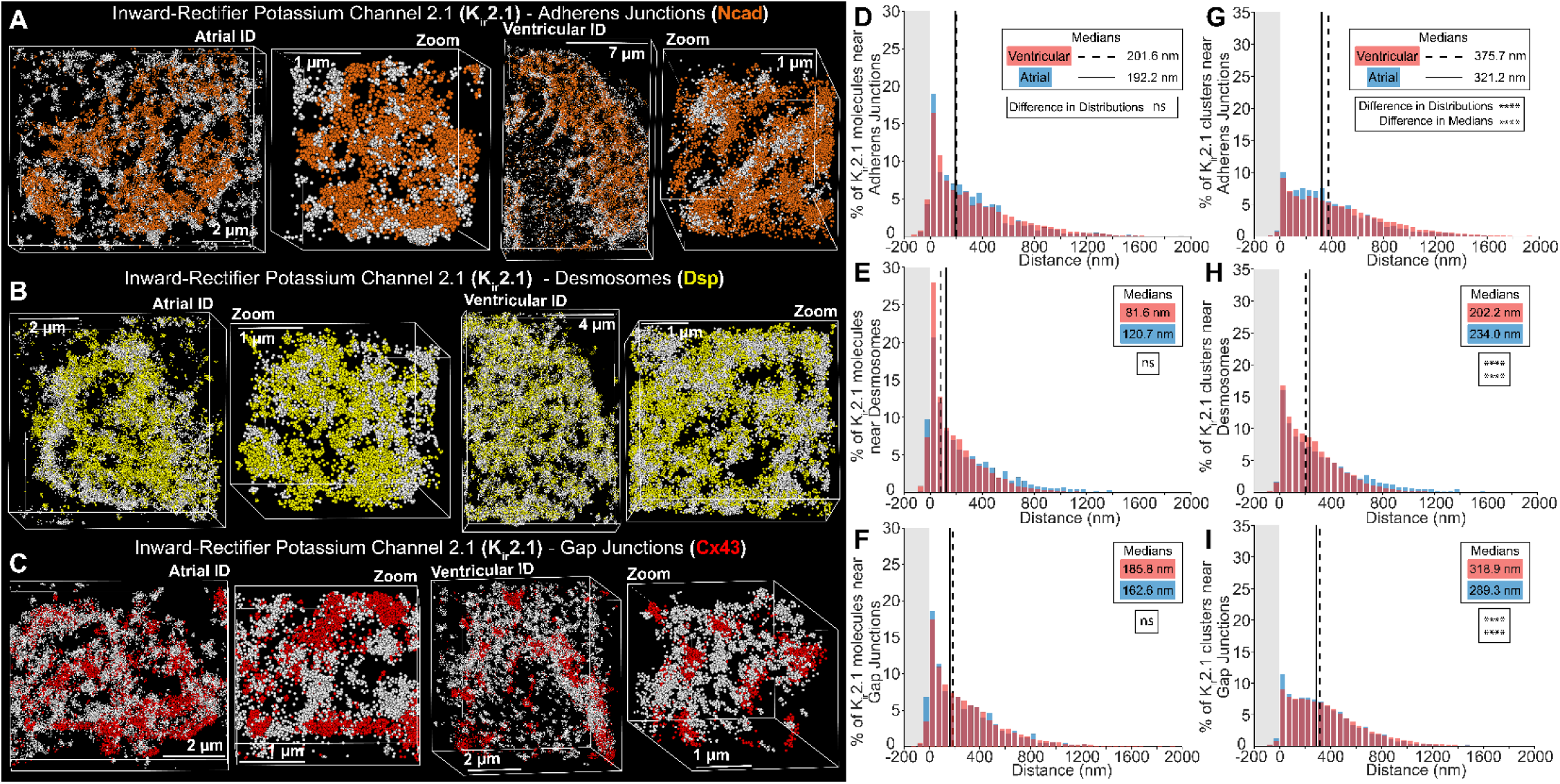
K_ir_2.1 channel distribution relative to junctional proteins. Representative STORM images of atrial and ventricular intercalated disc labeled for protein components of inward-rectifier potassium channel 2.1 (K_ir_2.1; white), gap junctions (Cx43; red), adherens junctions (Ncad; orange), and desmosomes (Dsp; yellow). **A)** inward-rectifier potassium channel 2.1 + adherens junctions, **B)** inward-rectifier potassium channel 2.1 + desmosomes, and **C)** inward-rectifier potassium channel 2.1 + gap junctions. Images were presented as whole intercalated disc and zoomed in views. Protein localization measured as **D-F)** % molecules and **G-I)** % clusters of protein localized relative to the co-labeled protein (n= 5[A], 5[V] images/heart from 3 hearts). Distributions represented in red for ventricular and blue for atrial measurements. Medians were represented as dashed line for ventricular and solid line for atrial. Differences in distributions and medians were statistically proven with two-sample Kolmogorov-Smirnov test and Wilcoxon signed-rank test, respectively (p > 0.05: ns, p ≤ 0.05: *, p ≤ 0.01: **, p ≤ 0.001: ***, p ≤ 0.0001: ****).

**Supplementary Figure 10.**
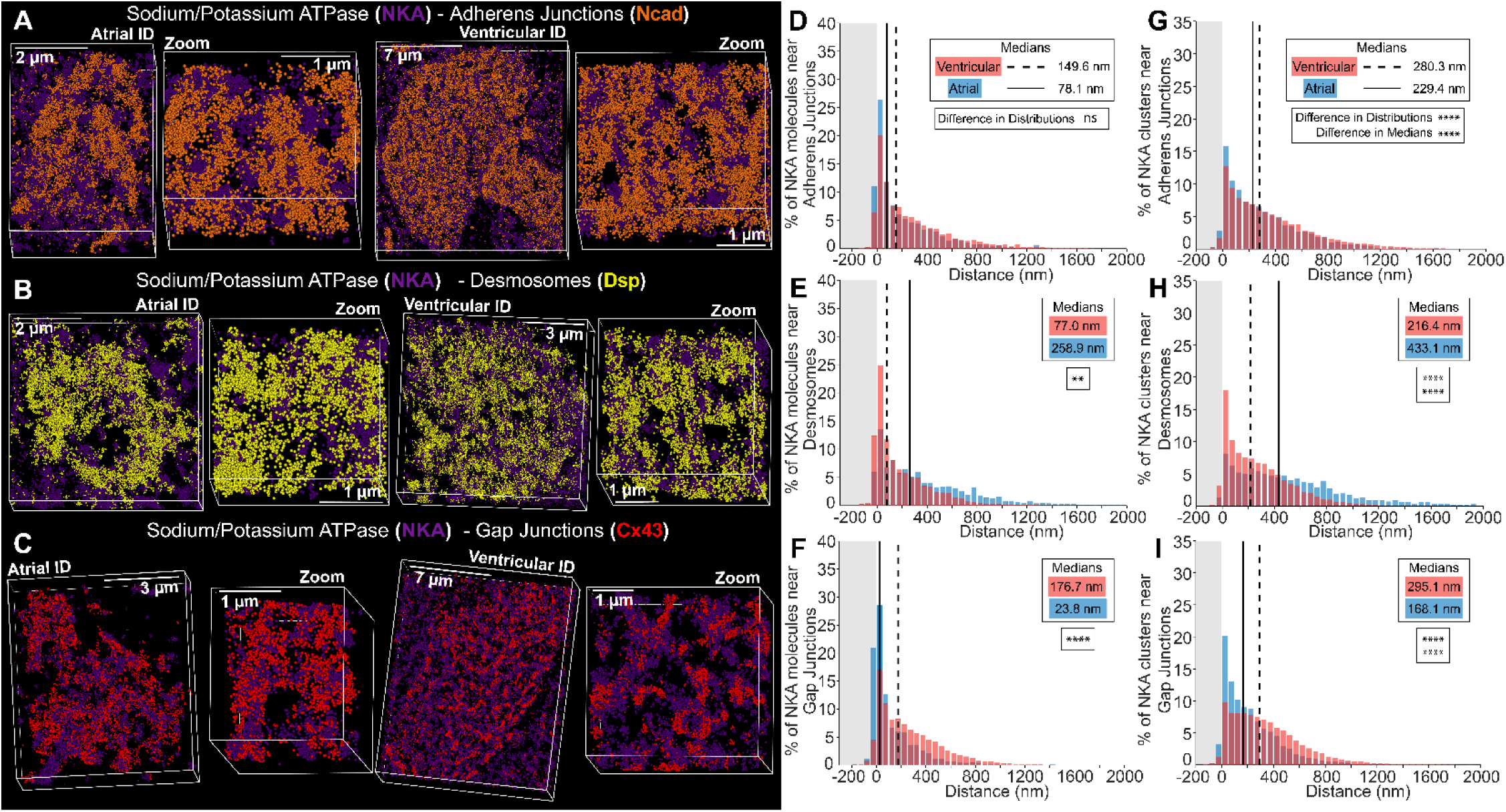
NKA channel distribution relative to junctional proteins. Representative STORM images of atrial and ventricular intercalated disc labeled for protein components of sodium potassium ATPase (NKA; purple), gap junctions (Cx43; red), adherens junctions (Ncad; orange), and desmosomes (Dsp; yellow). **A)** sodium potassium ATPase + adherens junctions, **B)** sodium potassium ATPase + desmosomes, and **C)** sodium potassium ATPase + gap junctions. Images were presented as whole intercalated disc and zoomed in views. Protein localization measured as **D-F)** % molecules and **G-I)** % clusters of protein localized relative to the co-labeled protein (n= 5[A], 5[V] images/heart from 3 hearts). Distributions represented in red for ventricular and blue for atrial measurements. Medians were represented as dashed line for ventricular and solid line for atrial. Differences in distributions and medians were statistically proven with two-sample Kolmogorov-Smirnov test and Wilcoxon signed-rank test, respectively (p > 0.05: ns, p ≤ 0.05: *, p ≤ 0.01: **, p ≤ 0.001: ***, p ≤ 0.0001: ****).

**Supplementary Figure 11.**
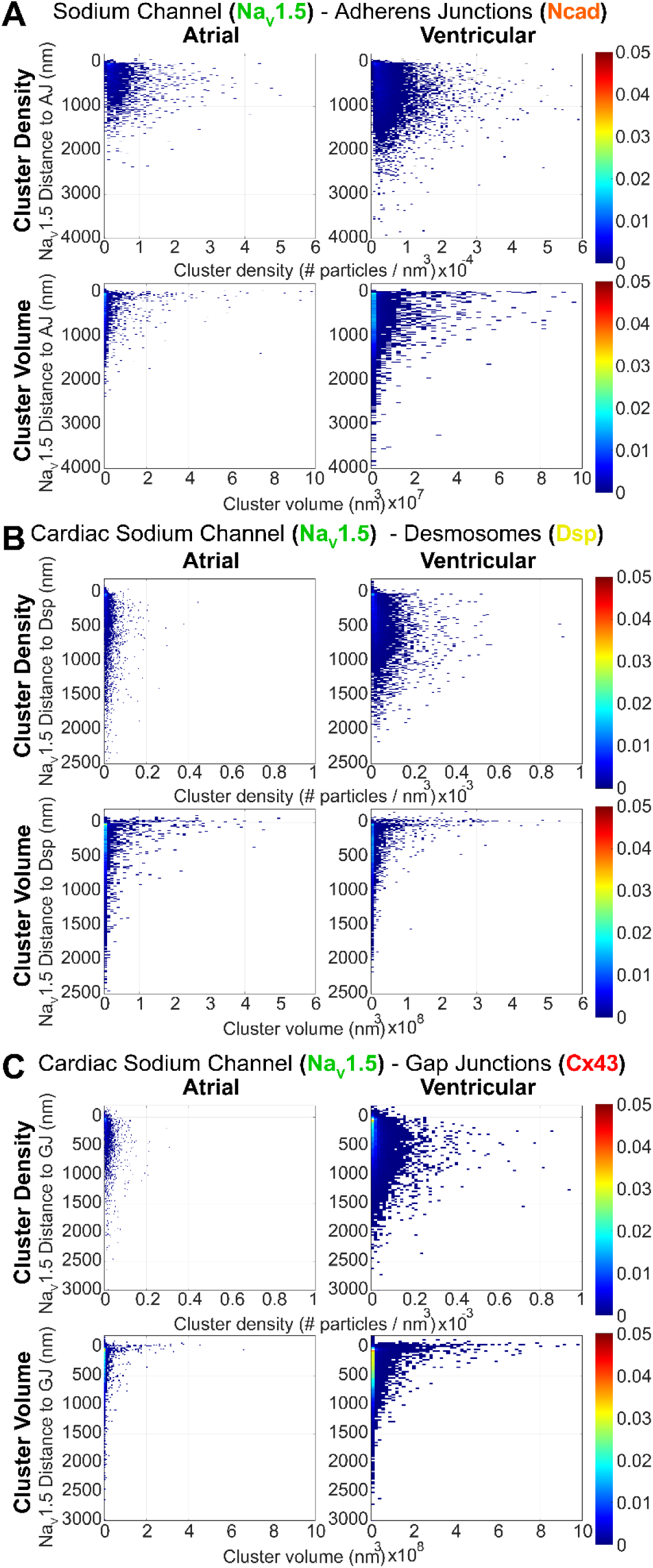
Cluster density and volume for Na_V_1.5 relative to junctional proteins. Bivariate histograms of cluster density and volume relative to distance from landmark proteins generated by STORM-based relative localization analysis (STORM-RLA). Analysis of Na_V_1.5 channel clustering: Na_V_1.5 relative to **A)** adherens junctions, **B)** desmosomes, and **C)** gap junctions.

**Supplementary Figure 12.**
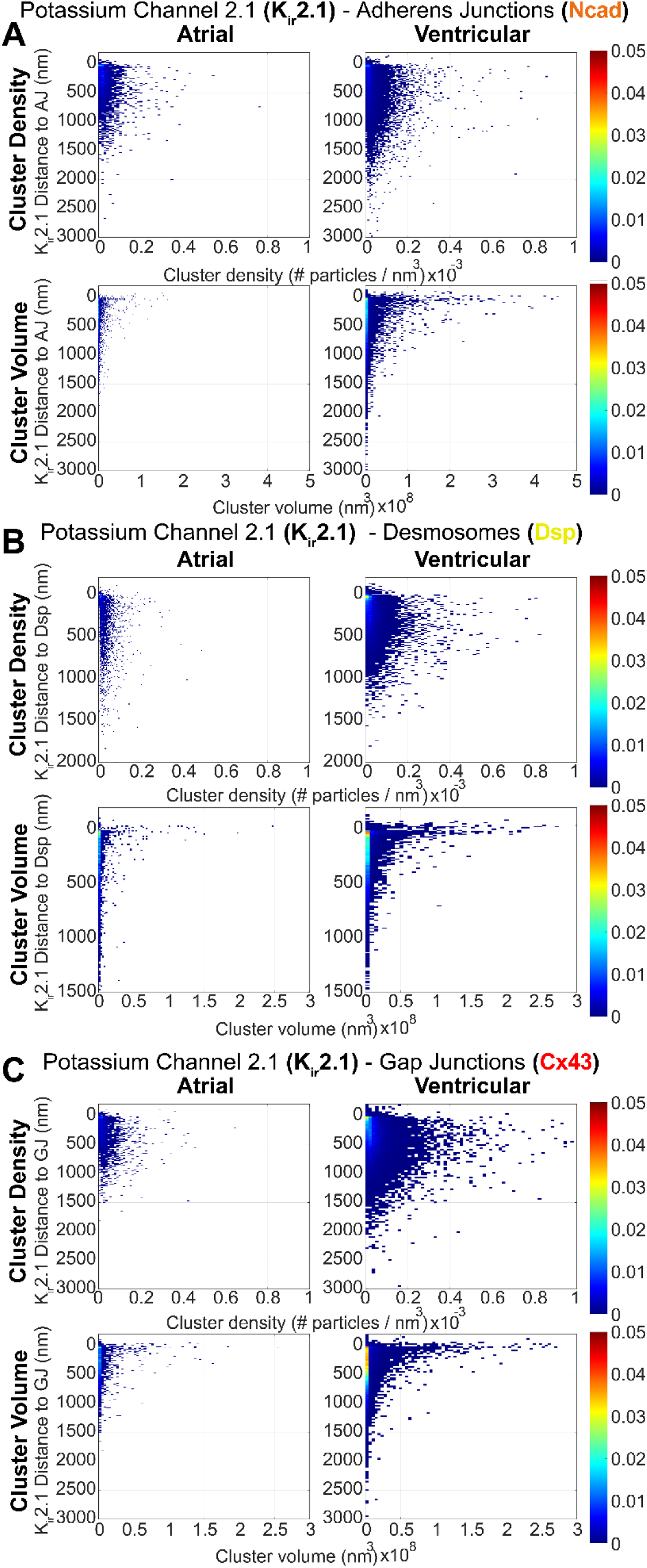
Cluster density and volume for K_ir_2.1 relative to junctional proteins. Bivariate histograms of cluster density and volume relative to distance from landmark proteins generated by STORM-based relative localization analysis (STORM-RLA). Analysis of K_ir_2.1 channel clustering: K_ir_2.1 relative to **A)** adherens junctions, **B)** desmosomes, and **C)** gap junctions.

**Supplementary Figure 13.**
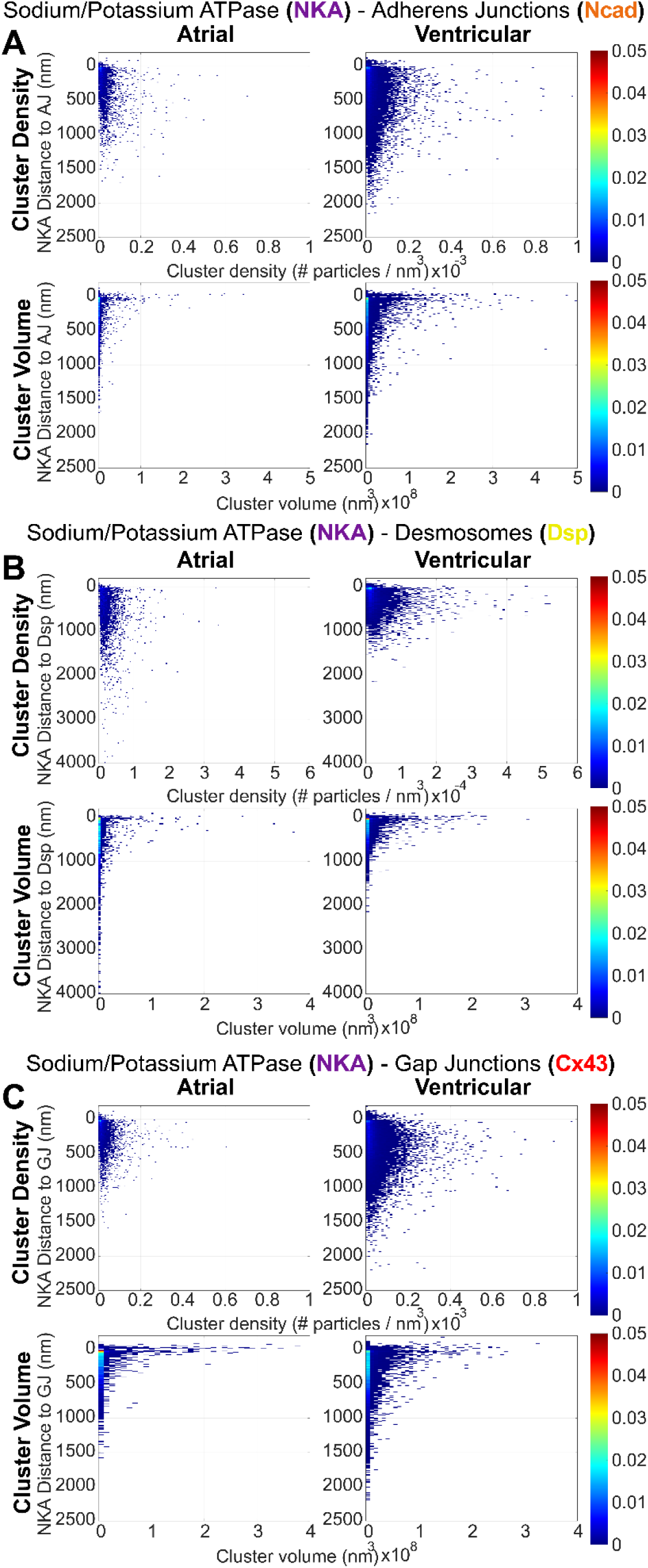
Cluster density and volume for NKA relative to junctional proteins. Bivariate histograms of cluster density and volume relative to distance from landmark proteins generated by STORM-based relative localization analysis (STORM-RLA). Analysis of NKA channel clustering: NKA relative to **A)** adherens junctions, **B)** desmosomes, and **C)** gap junctions.

**Supplementary Figure 14.**
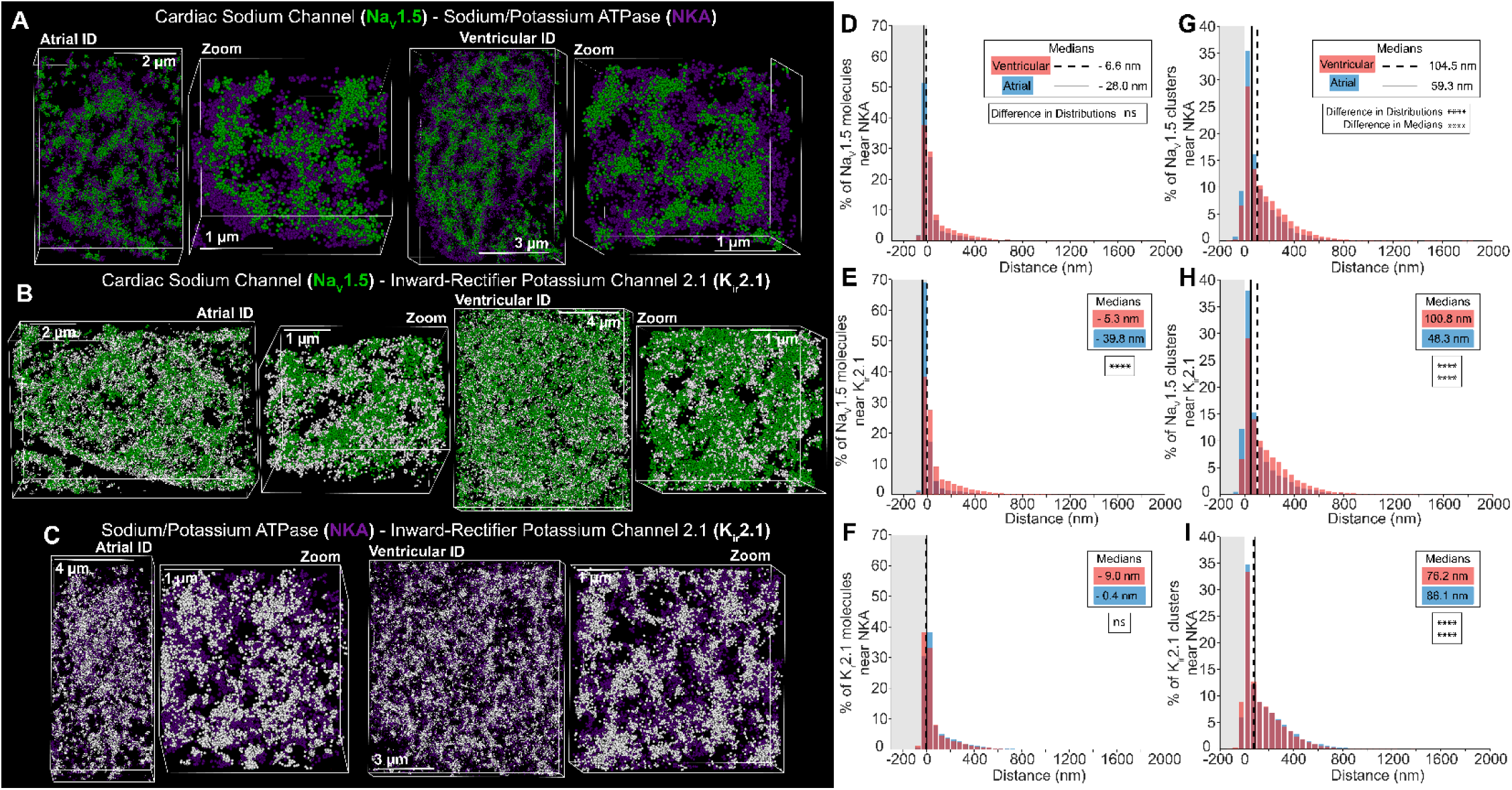
Distribution of electrogenic proteins relative to each other. Representative STORM images of atrial and ventricular intercalated disc labeled for protein components of cardiac isoform of the voltage-gated sodium channel (Na_V_1.5; green), inward-rectifier potassium channel (K_ir_2.1; white), and sodium potassium ATPase (NKA; purple). **A)** cardiac isoform of the voltage-gated sodium channel + sodium potassium ATPase, **B)** cardiac isoform of the voltage-gated sodium channel + inward-rectifier potassium channel, and **C)** sodium potassium ATPase + inward-rectifier potassium channel. Images were presented as whole intercalated disc and zoomed in views. Protein localization measured as **D-F)** % molecules and **G-I)** % clusters of protein localized relative to the co-labeled protein (n= 5[A], 5[V] images/heart from 3 hearts). Distributions represented in red for ventricular and blue for atrial measurements. Medians were represented as dashed line for ventricular and solid line for atrial. Differences in distributions and medians were statistically proven with two-sample Kolmogorov-Smirnov test and Wilcoxon signed-rank test, respectively (p > 0.05: ns, p ≤ 0.05: *, p ≤ 0.01: **, p ≤ 0.001: ***, p ≤ 0.0001: ****).

**Supplementary Figure 15.**
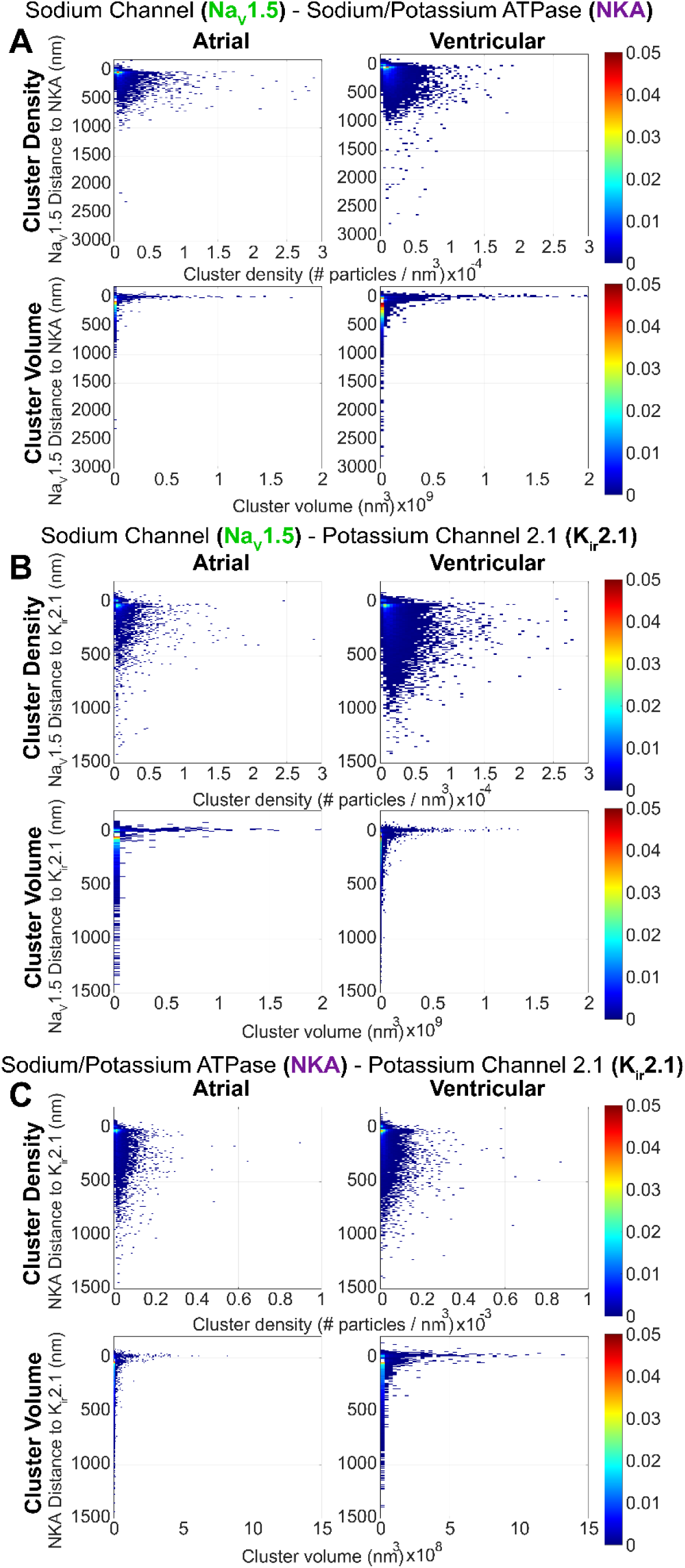
Cluster density and volume for electrogenic proteins relative to each other. Bivariate histograms of cluster density and volume relative to distance from landmark proteins generated by STORM-based relative localization analysis (STORM-RLA). Analysis of electrogenic channel clustering: **A)** Na_V_1.5 relative to NKA, **B)** Na_V_1.5 relative to K_ir_2.1, and **C)** NKA relative to K_ir_2.1.

**Supplementary Table 1.** Myocyte Structure and Intercalated Disc Ultrastructure Measurements. Cardiomyocyte size and intercalated disc ultrastructural properties are summarized by median, mean, standard error, number of measurements (n), and quartile range (Q1-Q3).

| Ultrastructure Organization Measurements |  |  |
| --- | --- | --- |
| Median \| Mean ± Standard Error (n) (Quartile range, Q1-Q3) | | |
| Cardiomyocyte Size |  |  |
|  | Atrial | Ventricular |
| Cardiomyocyte Length ($\mu\text{m}$) | 88.4 \| 88.8 ± 10.7 (151) (75.3-100.2) | 127.3 \| 128.6 ± 15.5 (150) (109.4-146.6) |
| Cardiomyocyte Width ($\mu\text{m}$) | 17.9 \| 18.9 ± 2.9 (151) (15.3-21.2) | 31.4 \| 33.2 ± 5.6 (150) (26.6-38.6) |
| Cardiomyocyte Depth ($\mu\text{m}$) | 8.9 \| 8.7 ± 1.6 (151) (6.8-9.8) | 11.4 \| 11.6 ± 1.4 (150) (9.8-13.2) |
| Cardiomyocyte Volume ($\mu\text{m}^3$) | 5,784.4 \| 6,170.6 ± 1,712.5 (151) (4,281.9-7,245.9) | 19,447.1 \| 20,761.9 ± 50,534.0 (150) (14,871.3-24,739.5) |
| Intercalated Disc Size |  |  |
|  | Atrial | Ventricular |
| Total Intercalated Disc Length ($\mu\text{m}$) | 4.9 \| 5.7 ± 1.8 (54) (3.5-7.5) | 9.0 \| 10.4 ± 2.8 (58) (6.8-12.5) |
| Intercalated Disc Volume ($\mu\text{m}^3$) | 266.3 \| 278.6 ± 70.8 (151) (201.8-312.4) | 1,055.8 \| 1,086.9 ± 253.0 (150) (802.8-1,294.6) |
| Plicate Ultrastructure Characteristics |  |  |
|  | Atrial | Ventricular |
| Plicate ID Fraction (%) | 57.90 \| 54.72 ± 10.39 (53) (41.42-67.78) | 62.53 \| 61.88 ± 7.50 (58) (54.02-70.51) |
| Plicate Length ($\mu\text{m}$) | 1.22 \| 1.35 ± 0.47 (174) (0.72-1.88) | 1.35 \| 1.76 ± 0.78 (247) (0.74-2.37) |
| Frequency (#waves/distance) | 2.23 \| 2.35 ± 0.50 (72) (1.72-2.91) | 2.47 \| 2.54 ± 0.47 (110) (1.90-3.03) |
| Amplitude ($\mu\text{m}$) | 0.06 \| 0.08 ± 0.03 (79) (0.04-0.08) | 0.16 \| 0.17 ± 0.04 (128) (0.12-0.23) |
| Plicate MJ membrane fraction (%) | 74.00 \| 69.76 ± 8.70 (57) (60.54-80.25) | 67.91 \| 68.30 ± 7.89 (52) (56.60-77.27) |
| Plicate MJ Length ($\mu\text{m}$) | 0.19 \| 0.26 ± 0.11 (282) (0.13-0.32) | 0.27 \| 0.33 ± 0.14 (329) (0.17-0.40) |
| Plicate GJ membrane fraction (%) | 12.59 \| 12.33 ± 3.97 (19) (7.50-15.50) | 8.82 \| 10.89 ± 3.91 (17) (5.44-15.43) |
| Plicate GJ Length ($\mu\text{m}$) | 0.12 \| 0.17 ± 0.10 (29) (0.09-0.20) | 0.18 \| 0.21 ± 0.08 (56) (0.14-0.24) |
| Intermembrane Distance Inside MJ (nm) | 13.43 \| 13.70 ± 1.60 (178) (11.54-15.22) | 16.88 \| 17.14 ± 1.96 (323) (14.67-19.32) |
| Intermembrane Distance 10 nm Outside MJ (nm) | 12.71 \| 13.18 ± 1.80 (178) (11.09-14.77) | 17.43 \| 17.59 ± 2.92 (241) (13.94-20.94) |
| Intermembrane Distance 30 nm Outside MJ (nm) | 11.92 \| 12.54 ± 2.11 (178) (9.95-14.28) | 16.39 \| 17.08 ± 3.57 (240) (12.32-20.38) |
| Interplicate Ultrastructure Characteristics |  |  |
|  | Atrial | Ventricular |
| Interplicate ID Fraction (%) | 42.10 \| 45.28 ± 10.39 (53) (32.22-58.58) | 36.72 \| 37.48 ± 7.39 (55) (27.50-44.76) |
| Interplicate Length ($\mu\text{m}$) | 1.23 \| 1.60 ± 0.74 (97) (0.84-1.69) | 1.34 \| 1.53 ± 0.51 (160) (0.92-1.94) |
| Interplicate MJ membrane fraction (%) | 10.00 \| 9.19 ± 2.78 (11) (4.98-12.19) | 5.93 \| 7.86 ± 3.06 (15) (4.46-10.11) |
| Interplicate MJ Length ($\mu\text{m}$) | 0.14 \| 0.16 ± 0.05 (57) (0.11-0.18) | 0.11 \| 0.13 ± 0.05 (24) (0.09-0.13) |
| Interplicate GJ membrane fraction (%) | 34.31 \| 39.68 ± 14.65 (40) (18.00-57.00) | 47.55 \| 46.43 ± 11.31 (72) (32.93-60.64) |
| Interplicate GJ Length ($\mu\text{m}$) | 0.33 \| 0.38 ± 0.14 (162) (0.21-0.47) | 0.38 \| 0.48 ± 0.22 (186) (0.25-0.59) |
| Intermembrane Distance 10 nm Outside GJ (nm) | 9.38 \| 9.65 ± 1.52 (58) (7.75-11.23) | 9.94 \| 10.55 ± 2.13 (142) (7.89-11.99) |
| Intermembrane Distance 30 nm Outside GJ (nm) | 11.50 \| 12.14 ± 2.40 (58) (9.82-12.67) | 12.13 \| 14.08 ± 3.51 (143) (9.63-16.68) |
| Intermembrane Distance 50 nm Outside GJ (nm) | 11.69 \| 12.90 ± 3.03 (58) (9.26-13.89) | 14.17 \| 15.91 ± 4.11 (140) (10.30-19.83) |

**Supplementary Table 2.**
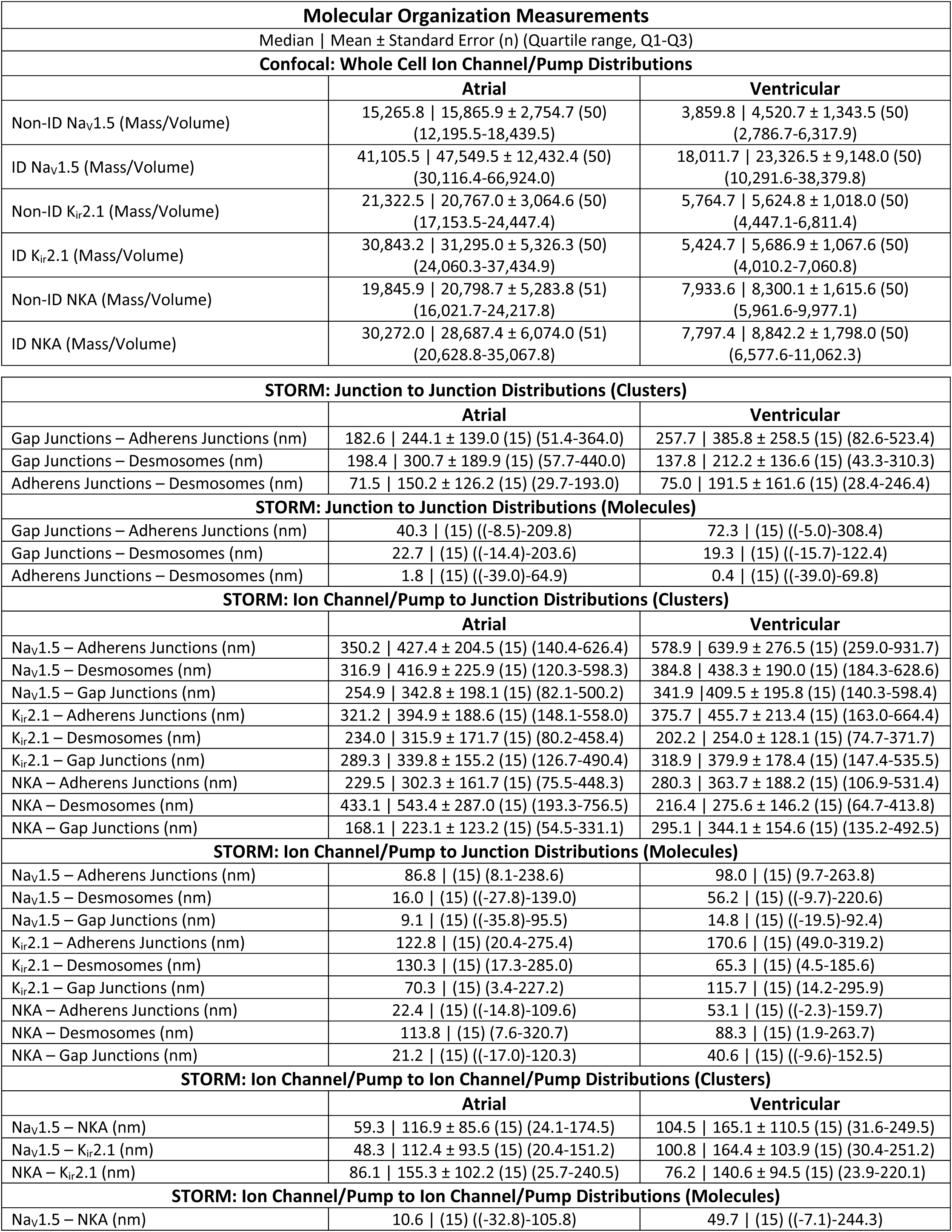
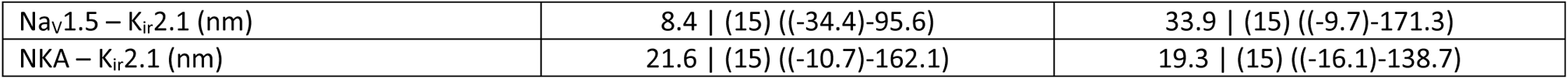
Molecular organization measurements. Measurements of molecular organization are summarized by median, mean, standard error, number of images (n), and quartile range (Q1-Q3).

**Supplementary Table 3.**
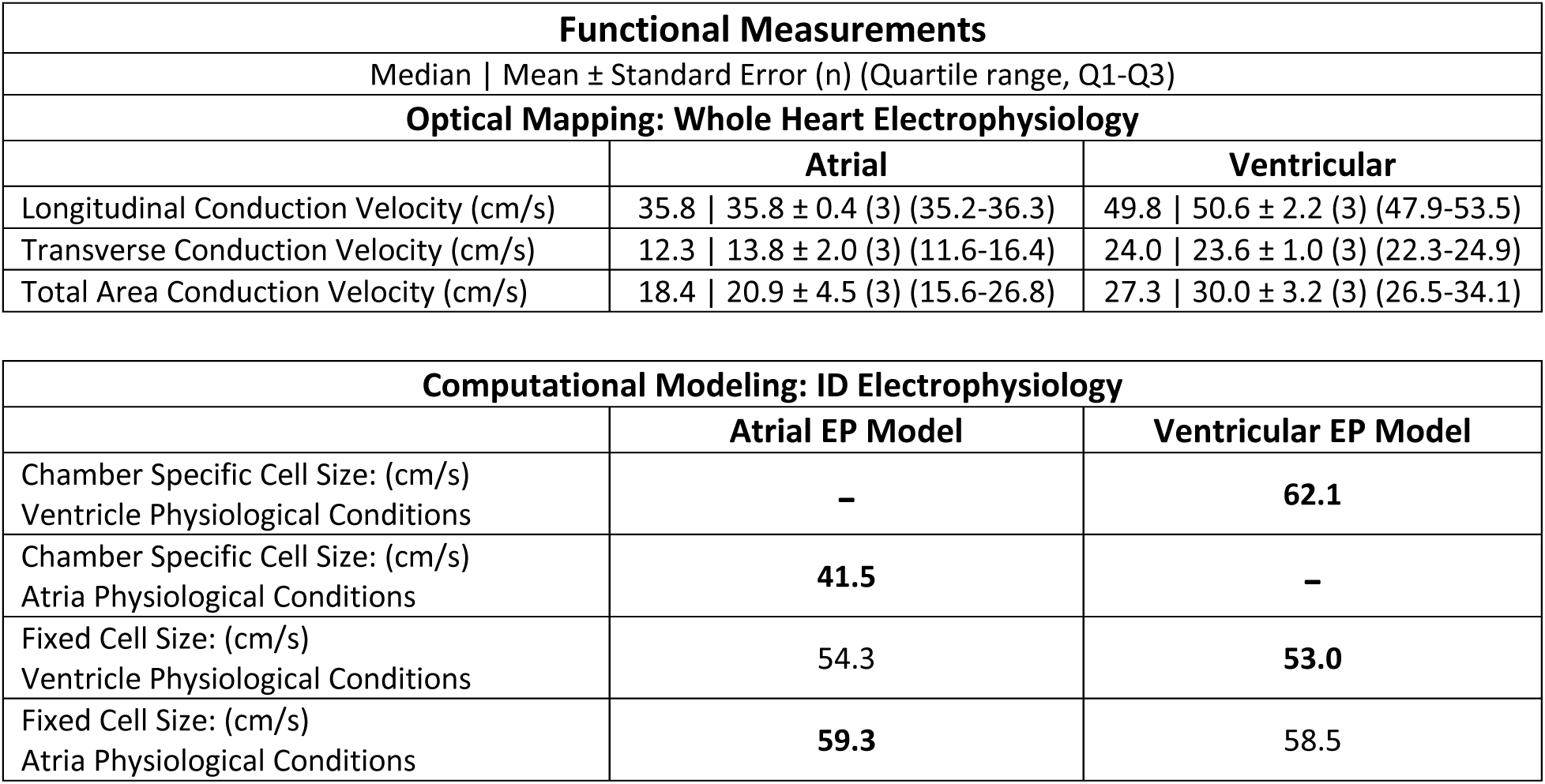
Functional measurements. Functional results are summarized by median, mean, standard error, number of images (n), and quartile range (Q1-Q3). Bolded numbers represent chamber-specific structures integrated with corresponding EP models (i.e. Atrial structure with atrial EP model).

